# Variable recruitment of distal tuft dendrites shapes new hippocampal place fields

**DOI:** 10.1101/2024.02.26.582144

**Authors:** Justin K. O’Hare, Jamie Wang, Margjele D. Shala, Franck Polleux, Attila Losonczy

## Abstract

Hippocampal pyramidal neurons support episodic memory by integrating complementary information streams into new ‘place fields’. Distal tuft dendrites are widely thought to initiate place field formation by locally generating prolonged, globally-spreading *Ca^2+^*spikes known as plateau potentials. However, the hitherto experimental inaccessibility of distal tuft dendrites in the hippocampus has rendered their *in vivo* function entirely unknown. Here we gained direct optical access to this elusive dendritic compartment. We report that distal tuft dendrites do not serve as the point of origin for place field-forming plateau potentials. Instead, the timing and extent of peri-formation distal tuft recruitment is variable and closely predicts multiple properties of resultant place fields. Therefore, distal tuft dendrites play a more powerful role in hippocampal feature selectivity than simply initiating place field formation. Moreover, place field formation is not accompanied by global *Ca^2+^*influx as previously thought. In addition to shaping new somatic place fields, distal tuft dendrites possess their own local place fields. Tuft place fields are back-shifted relative to that of their soma and appear to maintain somatic place fields via post-formation plateau potentials. Through direct *in vivo* observation, we provide a revised dendritic basis for hippocampal feature selectivity during navigational learning.

## INTRODUCTION

The central nervous system supports learning and memory by accomplishing two core tasks: (1) integrating sensory information to faithfully represent the external world and (2) updating how it processes this information such that future computations better promote an organism’s overall fitness. Pyramidal neurons (PNs) bear much of this responsibility as they comprise the main source of excitatory drive in the cerebral cortex^1, 2^. As such, PNs unite multiple levels of brain function to carry out complex computations. A single PN receives many thousands of synaptic inputs across its dendritic arbor^3, 4^. These inputs typically originate from multiple presynaptic circuits that broadcast complementary streams of information to distinct dendritic compartments^5, 6^. In turn, dendritic compartments can locally and nonlinearly process synaptic inputs^7-12^ according to bespoke integrative rules^13-17^ while local plasticity mechanisms update these rules with experience^18-22^. This nexus between subcellular and systems levels of brain function endows a single PN with considerable computational capacity^23-27^ but is exceedingly difficult to interrogate experimentally. Therefore, one of the most central questions in neuroscience is how PNs integrate their diverse synaptic inputs to drive feature-selective somatic action potential (AP) firing underlying behavioral adaptation^28, 29^.

PNs in hippocampal area CA1 represent a key cellular substrate for learning and memory and exhibit a striking form of feature selectivity in the form of receptive fields known as ‘place fields’ (PFs). PFs emerge from experience to conjunctively encode context-specific sensory features^30, 31^ and, as a result, are spatially tuned to specific locations within an animal’s environment^32^. PFs support spatial navigation in rodents by forming flexible ‘cognitive maps’^32-34^ and analogous receptive fields are believed to support episodic learning and memory in humans^35-38^. New PFs emerge from experience via a recently described synaptic plasticity mechanism known as behavioral timescale synaptic plasticity (BTSP) ^39^. BTSP is unique in its ability to bind together pre- and postsynaptic activity over behaviorally relevant timescales after a single pairing. For this reason, much recent effort has been dedicated to uncovering its still-elusive circuit^40, 41^, cellular^20, 41-43^, and molecular^44-46^ mechanisms.

Distal tuft dendrites are thought to initiate PF formation. Specifically, they are thought to convert multiplexed sensory and spatial signals from the entorhinal cortex (EC) ^47-56^ into an instructive signal in the form of a dendritic plateau potential^39, 42, 57^. According to this model, plateau potentials depolarize the entire PN such that any dendritic spines receiving excitatory input within the seconds-long BTSP association window may undergo plasticity. This model is supported by causal manipulations of EC inputs to CA1^40, 57, 58^, the re-weighting of more-proximal synapses during PF formation^20, 41, 43^, and the ability of distal dendrites to drive global depolarization and promote synaptic plasticity^59-62^ *in vitro*. However, distal tuft dendrites have hitherto proven experimentally inaccessible in CA1 PNs due to their location at the deepest reaches of the apical arbor. The inability to directly monitor this elusive dendritic compartment has significantly impaired our ability to test its role in the emergence of hippocampal feature selectivity.

Here we gained direct, simultaneous access to distal tuft dendrites and their soma in single CA1 PNs *in vivo*. We addressed, for the first time, three broad questions: (1) How does distal tuft dendritic activity inform somatic activity across behavioral states? (2) What experiential features do distal tuft dendrites encode? (3) What roles do distal tuft dendrites play in the emergence of new PFs during spatial navigation? We leveraged single-cell DNA electroporation, multi-plane two-photon *Ca^2+^* imaging, and a virtual reality-based serial ‘teleportation’ paradigm to uncover a remarkably broad repertoire of previously unappreciated distal dendritic functions that address each of these questions.

## RESULTS

### Monitoring somatic and distal tuft dendritic dynamics in single CA1 PNs *in vivo*

We sought to investigate the role of distal tuft dendrites in shaping somatic activity and forming new PFs. To do so required simultaneous access to the soma and distal tuft dendrites of CA1 PNs. However, these dendrites have not been previously imaged in CA1 PNs *in vivo* due to their depth and overlap with neighboring dendritic arbors. We first overcame these biological constraints by expressing DNA plasmids encoding the red-shifted *Ca^2+^* indicator XCaMP-R^63^ in individual CA1 PNs using *in vivo* single-cell electroporation (SCE) ^20, 64-66^ (Figure 1A). The use of a red-shifted sensor mitigated depth-dependent light scattering from biological tissue while SCE provided single-neuron sparsity. We then gained simultaneous access to the soma of imaged distal tuft dendrites by using a piezoelectric device to rapidly actuate a 40x objective lens between distant focal planes (Figure 1B). To relate somatic and distal dendritic dynamics to animal behavior and PFs, we imaged electroporated CA1 PNs while mice navigated a 3-m virtual linear track^67^ for randomly located water rewards (Figure 1, C & D; Figure S1). SCE permitted unambiguous allocation of imaged dendrites to their parent soma (Figure 1, E & F).

**Figure 1.**
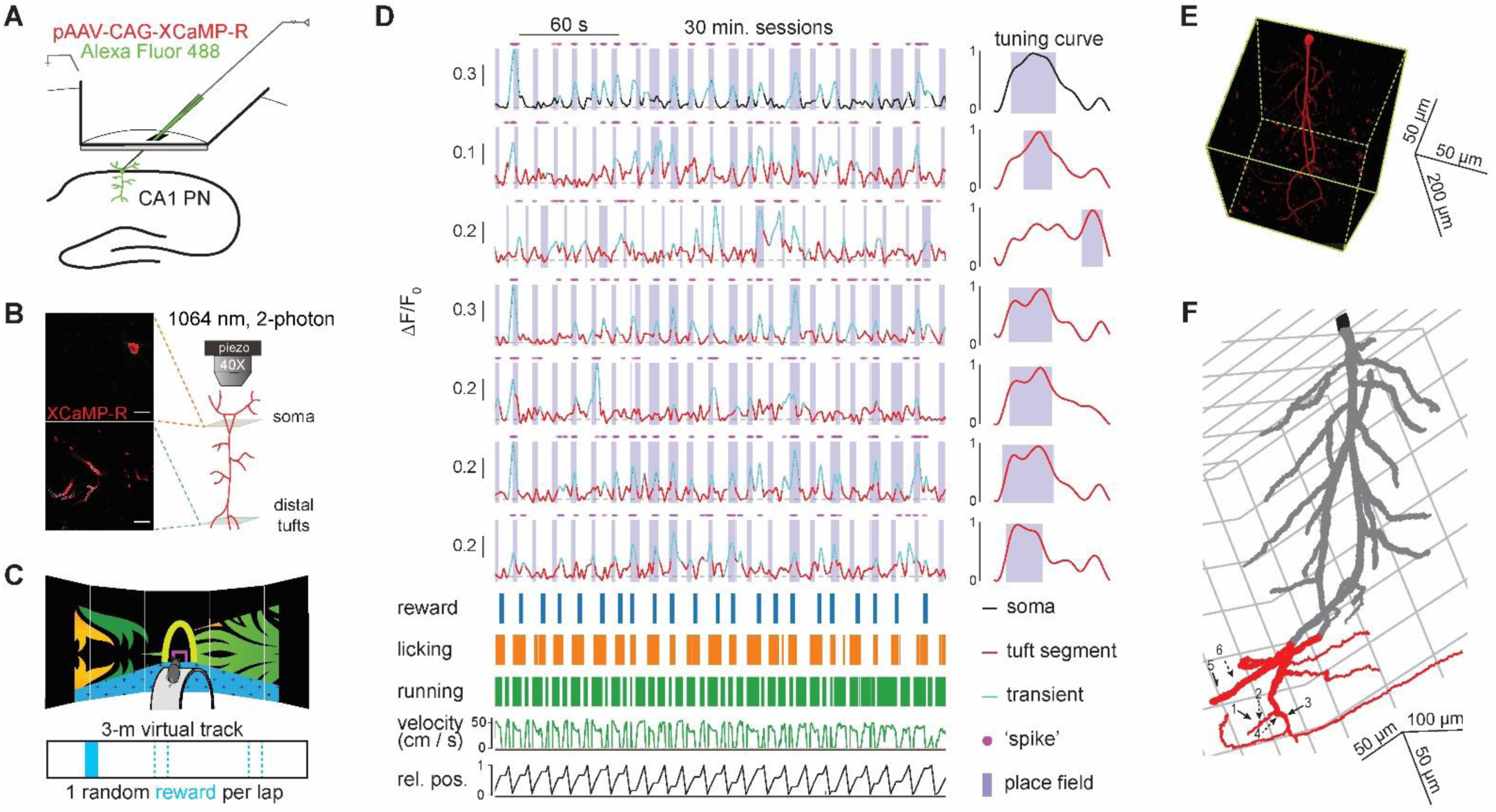
Simultaneously monitoring somatic and distal tuft dendritic dynamics in isolated CA1 PNs during virtual navigation. (**A**) Plasmid DNA encoding the red-shifted *Ca^2+^*indicator XCaMP-R was delivered to individual CA1 PNs in anesthetized adult mice. Alexa Fluor 488 was visualized by 2-photon microscopy to guide pipet descent. (**B**) XCaMP-R was simultaneously imaged in somatic and distal tuft dendritic imaging planes using a 1070 nm fixed-wavelength 2-photon laser and a piezoelectric device that rapidly toggled a 40x water immersion objective lens. Representative motion-corrected, time-averaged images are shown for each focal plane (scale bar: 20 µm). Images were cropped to emphasize regions of interest (white outlines). (**C**) Mice navigated a 3-m virtual linear track for randomly delivered water rewards (1 / lap). (**D**) Summary plot depicting somatic and dendritic signals with time-locked behavioral variables. (**E**) Representative *in vivo* volumetric scan of same CA1 PN shown in panels (B) and (D). Image stack was processed to maintain consistent contrast across depths. Basal dendrites dorsal to the soma not included. (**F**) Reconstruction of CA1 PN shown in (E), color-coded by compartment (soma: black, apical trunk and radial oblique dendrites: grey, distal tuft dendrites: red). Numbered arrows indicate imaged dendrites shown in same order as in (D). Dashed lines indicate dendrites that are occluded in 3D reconstruction.

### CA1 PN soma distal tuft dendrites are most strongly isolated from their soma during locomotion

The degree of functional autonomy that distal tuft dendrites possess relative to their soma carries major implications regarding their function, including the extent to which they filter EC inputs as well as their role in somatic PF formation. CA1 PN distal tuft dendrites are located > 300 microns from their soma with potentially limited influence on somatic AP firing due to passive^68, 69^ filtering properties of dendrites as well as voltage-dependent leak currents^70-73^. On the other hand, CA1 PN tuft dendrites can drive robust somatic AP firing under certain conditions including, but not limited to, rebound spiking from dendritic inhibition^74-76^, temporally correlated input to distal and proximal dendrites^60, 61, 77^, and somatic depolarization^73^. While these *in vitro* and computational studies have proven invaluable in appreciating the complexity of dendritic computation in the hippocampus, they have also highlighted the challenges inherent in predicting how tuft dendrites might influence their soma *in vivo* while sensory experience and animal behavior continuously and variably modulate each of the aforementioned determinants of tuft-soma crosstalk.

To gain insight into the functional properties of distal tuft dendrites, and garner clues as to how they might support higher-level cellular processes such as PF formation, we first examined basic *Ca^2+^* transient properties of these dendrites relative to their soma. Tuft dendrites possessed shorter *Ca^2+^* transient waveforms than their soma while both compartments displayed strong positive modulation by locomotion (Figure S2, A-C). Tuft dendrites displayed a high degree of autonomous activity, with many *Ca^2+^* transients lacking a contemporaneous somatic transient (‘isolated’ transients) or even a contemporaneous tuft transient (Figure 2A). Greater than one third of tuft *Ca^2+^* transients were isolated during running periods (med. = 0.37, IQR = 0.18) and, counter to our expectation, this fraction was reduced on average during periods of immobility (med. = 0.29, IQR = 0.31) (Figure 2B).

**Figure 2.**
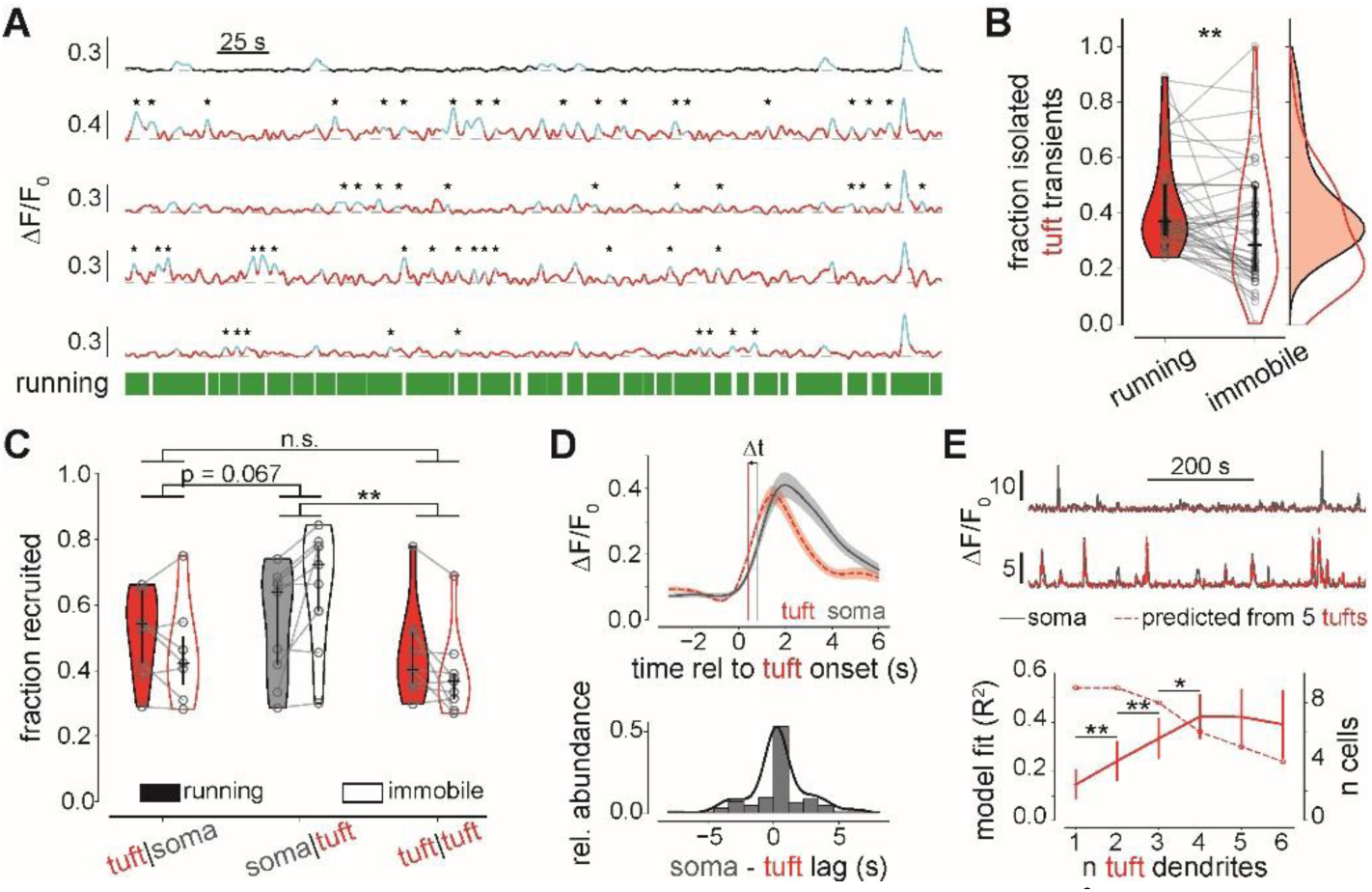
Bidirectional and state-dependent compartmentalization between CA1 PN soma and distal tuft dendrites. (**A**) Soma (black) and tuft (red) traces from an example CA1 PN. *Ca^2+^*transients are overlaid in cyan. Asterisks indicate isolated tuft transients. Binary running frames (green) indicate velocity ≥ 1 cm / s. (**B)** Fractions of tuft *Ca^2+^* transients that were isolated, i.e. did not propagate to soma, during running and immobile states. Gaussian kernel density estimates shown on right. (**C**) Cross- compartment conditional analysis quantifying the fraction of times compartment ‘a’ was recruited when compartment ‘b’ fired (‘a|b’). Values were averaged within-cell. Linear mixed effects model, compartments effect: *F*_(1.42, 32.70)_ = 6.50, *P* < 0.01; running effect: *F*_(1, 46)_ = 0.022, *P* > 0.05. Post hoc *t*-test results shown. (**D**) Top: Mean somatic *Ca^2+^* transient waveform triggered on tuft dendrite transient onsets. Vertical lines indicate timestamps for half-maximum amplitudes used for lag calculation. Bottom: Histogram of soma-tuft lags with Gaussian kernel density estimate. (**E**) Top: Example tuft-based predictions of somatic *ΔF/F_0_* (red) illustrating range of model performance. True somatic *ΔF/F_0_* traces are shown in black. Predictions are each based on 5 tuft dendrites for ease of comparison. Bottom: Model performance as a function of the number of dendrites used for model training. Linear mixed effects model: *F*_(0.83, 4.46)_ = 11.90, *P* < 0.05. Fisher’s LSD shown for *N* vs. *N* – 1 comparisons up to *N* = 6. Dashed line corresponds to second y-axis and indicates the number of cells for which as least *N* tuft dendrites (x-axis) were imaged. Violin plots depict full data range with lines at medians and error bars spanning first to third quartiles. \**P* < 0.05, \*\**P* < 0.01. See table S1 for sample sizes and table S2 for additional statistical details.

We next asked whether soma-tuft compartmentalization might be hysteretic, i.e. biased toward propagation in a particular direction. Our temporal resolution did not permit direct observation of event propagation. However, events that occur in tuft dendrites but not their soma can be reasonably interpreted as failures of centripetal (i.e. ‘forward’) propagation and vice-versa. We thus employed a cross-compartment conditional analysis to assess the likelihoods that tuft events accompany a somatic event (‘tuft|soma’), a somatic event accompanies a tuft event (‘soma|tuft’), and tuft events accompany a tuft event (‘tuft|tuft’). This analysis revealed strong overall compartmentalization both between and within compartments. Centripetal propagation from tufts to soma was stronger than propagation between individual tuft segments and non-significantly (*P* = 0.067) stronger than centrifugal propagation from soma to tufts (Figure 2C). Consistently, somatic *Ca^2+^* transients most commonly lagged those of tuft dendrites (median = 0.20 s, IQR = 0.60 s) (Figure 2D). This measurement corresponded to the limit of our temporal resolution set by image acquisition rate across distant focal planes (5-6 Hz). Isolated tuft *Ca^2+^*transients displayed higher amplitude, duration, and overall tuft recruitment than those coactive with the soma (Figure S2, D-F). We note that cross-compartmental relationships in *Ca^2+^* flux may differ from those of voltage due to a number of reasons, including divergent *Ca^2+^* handing mechanisms. Nonetheless, the marked disparities between isolated and global *Ca^2+^* events, along with cross-compartment conditional analyses in Figure 2C, indicate that (1) backpropagating somatic APs typically drive voltage-dependent *Ca^2+^* influx in only a subset of tuft dendrites and (2) even large, supralinear events recruiting many tuft dendrites can fail to drive detectable somatic *Ca^2+^* transients *in vivo*.

Given the prevalence and prominence of isolated events in tuft dendrites, we next asked how much information is shared between tuft dendrites and their connected soma. To this end, we trained a double-cross-validated Ridge regression model to predict somatic XCaMP-R signals (*ΔF/F_0_*) from random combinations of 1 to 6 connected tuft dendrites (see supplementary materials and methods). Tuft predictive power linearly summed up to the inclusion of 4 dendrites, at which point model performance plateaued at *R^2^*= 0.42 ± 0.092 (Figure 2E). We draw two conclusions from this result. First, the modest peak model performance indicates that the overall amount of information shared between these distant cellular compartments is limited. Second, the fact that model performance saturated with only 4 tuft dendrites indicates that the information that is shared between compartments is relatively low-dimensional. In summary, local tuft dynamics, while clearly robust, appear to have limited influence on somatic action potential firing during ordinary foraging behavior in a familiar environment.

### Robust detection of somatic place field formation and dendritic plateau potentials *in vivo*

The high degree of compartmentalization we observed in CA1 PN distal tuft dendrites is amenable to their presumed role in PF formation. While these dendrites exert modest influence on somatic AP firing under basal conditions (Figure 2, B & C), they can generate robust isolated events (Figure S2, D-F) which, if properly timed with input to proximal apical dendrites, may drive dendritic plateau potentials to form new PFs. In the absence of an experimental approach to directly monitor plateau potentials in CA1 PN dendrites, previous *in vivo* studies have inferred dendritic plateau potentials from their somatic vestige known as the after-depolarizing potential (ADP) ^39, 42, 57, 78^. Therefore, it remains unclear where plateau potentials originate and how globally they spread throughout the dendritic arbor *in vivo*, i.e. what fraction of eligible dendritic spines will undergo synaptic plasticity during PF formation. These unanswered questions are critical to understanding cellular mechanisms of memory formation. To probe the role of tuft-originating plateau potentials in PF formation, we sought to (1) reliably elicit spontaneous somatic PF formation events, (2) assess tuft *Ca^2+^* dynamics at the time of those events, and (3) disambiguate between ordinary tuft *Ca^2+^*transients and those driven by dendritic plateau potentials.

While optogenetic stimulation and somatic current injection protocols have recently emerged as a mean to induce PF formation^20, 39, 41-43, 66, 79^ we required an approach that would preserve the *in situ* role of distal tuft dendrites in PF formation. Exposure to novel contexts has been shown to promote PF formation in area CA1 PNs^80-82^ in the absence of neural stimulation. We thus leveraged a virtual context switching task as a causal tool to evoke spontaneous PF formation. Mice were serially ‘teleported’ every 12-15 laps across four novel, 3-m linear virtual tracks. In imaging sessions using fixed-location water reward, mice rapidly learned to slow down and lick as they approach reward zones (Figure S3). Mice then revisited the same contexts (‘familiar’) for the remainder of each 30-minute imaging session (Figure 3A; see supplementary materials and methods).

**Figure 3.**
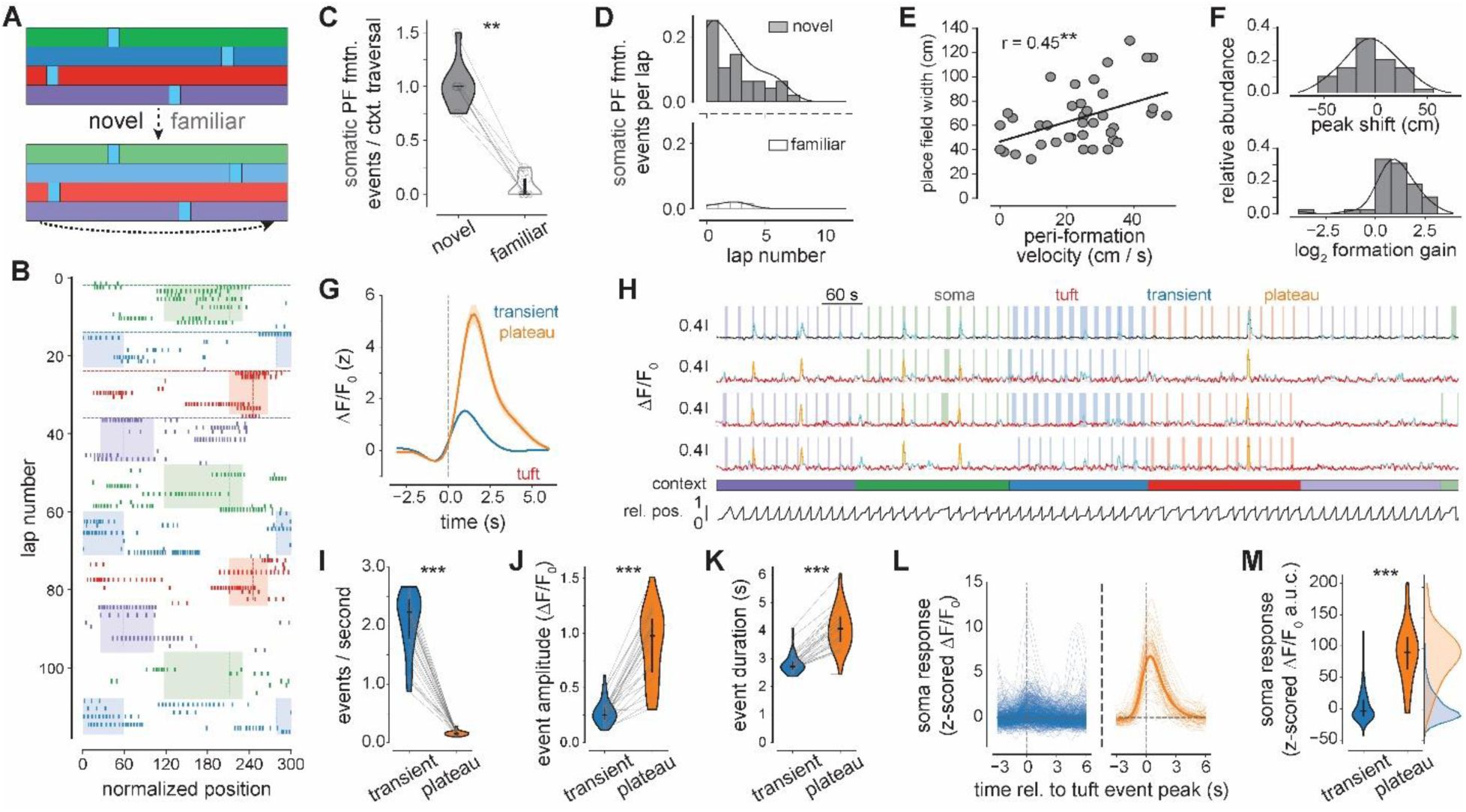
Detection and characterization of PF formation events and plateau potentials. (**A**) Schematic of virtual teleportation paradigm. All contexts were novel upon first exposure. In 6/8 experiments, reward was positioned at a unique fixed location in each context to promote attention to visual cues. (**B**) Example raster plot of somatic activity, color-coded by context. Tick marks represent deconvolved events from *ΔF/F_0_* signal which were used for spatial tuning analyses. Each row represents one lap. Shaded regions denote *de novo* PFs, horizontal dashed lines indicate PF formation laps, and vertical dashed lines indicate animal position at the moment of PF formation. (**C**) Somatic PF formation events per traversal of a novel or familiar context. (**D-F**) Hallmarks of BTSP-driven PF formation^20, 39, 43, 82^ including a bias for PFs to form early in novel contexts (D), a correlation between peri-formation running velocity and resultant PF width (E), a tendency for PFs to appear back-shifted in space relative to animal position at time of plateau potential (F, top), and stronger somatic activity during PF formation relative to subsequent PF traversals (F, bottom). (**G**) Mean event *Ca^2+^* waveforms for tuft transients (blue) and plateau potentials (orange). (**H**) Example somatic (black) and tuft (red) *Ca^2+^*traces showing both transients and plateau potentials with context and relative animal position at bottom. Shaded regions indicate PF traversals. (**I-K**) Comparison of basic *Ca^2+^* event properties between ordinary tuft *Ca^2+^* transients and plateau potentials including frequency (I), amplitude (J), and duration (K). (**L**) Somatic responses to ordinary *Ca^2+^*transients (blue, left) and plateau potentials (orange, right) detected from mean *ΔF/F_0_* signal of tuft dendrites imaged for each CA1 PN. (**M**) Quantification of somatic responses in (L). Violin plots depict full data range with lines at medians and error bars spanning first to third quartiles. \*\**P* < 0.01, \*\*\**P* < 0.001. See table S1 for sample sizes and table S2 for additional statistical details.

We then developed a new approach to detect and precisely timestamp PF formation events based on *Ca^2+^* signals (see supplementary materials and methods) and identified a remarkable 39 *de novo* somatic PF formation events from 30-min recordings of just 9 CA1 PNs (Figure 3, B & C; Figure S4). These PF formation events displayed distinctive hallmarks of BTSP^20, 39, 82^ including a strong association with novel context exposure, a correlation between peri-formation animal running speed and resultant PF width, a tendency for a new PF to emerge backward in space relative to the animal’s location at the time of PF formation, and stronger than usual peri-formation activity levels (Figure 3, C-F). *De novo* PFs were highly stable across laps as well as repeated traversals of the virtual contexts in which they emerged, despite interleaving exposure to 3 distinct contexts (Figure S5). Therefore, our virtual teleporting task served as a causal tool to reliably evoke PF formation while preserving the *in situ* role of tuft dendrites in this process.

To gain insight specifically into the role of tuft-driven plateau potentials in PF formation, we devised an unsupervised machine learning approach to factorize and cluster tuft *Ca^2+^* transient waveforms into two groups (Figure S6, A-C). Imposing zero assumptions regarding plateau potential *Ca^2+^*signals, we identified two distinct activity modes of tuft dendrites (Figure 3G). We then trained a double-cross-validated ensemble of support vector classifiers to correctly predict clustering labels of individual *Ca^2+^* events with an accuracy of 99.80% (Figure S6D). Next, we reran our *Ca^2+^* transient detection pipeline using template waveforms from both clusters along with the trained classifier to label events as ordinary *Ca^2+^* transients or putative plateau potentials (Figure S6E; Figure 3H). Finally, because plateau potentials are considered globally depolarizing cellular events, we applied a winner-take-all rule whereby all imaged tuft dendrites were required to fire an event with at least 50% of those events classified as a plateau. If this condition was met, the classification was applied to all tuft dendrites; otherwise, the event was labeled as an ordinary *Ca^2+^* transient. Putative plateau potentials possessed markedly distinct waveforms (Figure S6E; Figure 3, G & H) that were large, long, and rare relative to *Ca^2+^* transients (∼1/12^th^ the frequency) (Figure 3, I-K). Consistent with *in vitro* recordings^57, 61^, putative tuft plateau potentials elicited significantly stronger somatic responses (Figure 3, L & M) than did *Ca^2+^* transients. Therefore, while our identified plateau potentials are necessarily putative in nature, we take their numerous correspondences to *in vitro* measurements, as well as the unbiased manner in which they were identified, as a strong indicator of external validity.

### Distal tuft dendrites are reliably but variably recruited during somatic place field formation

Having developed approaches to promote and detect spontaneous somatic PF formation, and to detect plateau potentials, we first asked a simple question: what do distal tuft dendrites do while a CA1 PN forms a new somatic PF? Based on numerous lines of converging evidence, we hypothesized that distal tuft dendrites would generate plateau potentials to drive global dendritic *Ca^2+^* influx. Consistent with this hypothesis, somatic PF formation-triggered tuft *Ca^2+^* dynamics revealed that CA1 PN tuft dendrites are robustly activated during PF formation (Figure 4A). Distal tuft dendrites displayed larger than normal *Ca^2+^* events during PF formation (Figure 4B) and with peri-formation activation levels that were tightly coupled to those of their soma (Figure 4C). However, we noted two apparent discrepancies between peri-formation tuft activity and their purported role of initiating PF formation via locally generated plateaus. First, the variability of tuft peri-formation activity (Figure 4, B & C) appeared high given the stereotyped depolarizations observed during plateau potential-mediated PF formation in CA1 PNs^57^. Second, whereas novel context exposure robustly promoted somatic PF formation in this study (Figure 3, C & D) and in previous studies^80-82^, it did not influence tuft plateau frequency (Figure S7). In fact, novelty was associated with a tonic decoupling of tuft dendrites from their soma and an increased rate of isolated tuft *Ca^2+^* transients (Figure S8, A-C). Isolated tuft events were biased toward somatic PFs and predominantly occurred during post-formation laps. Therefore, isolated tuft events did not appear to predict future PF formation events (Figure S8D) as has been observed in CA1 PN basal dendrites^80^. Taken together, these observations affirmed the engagement of tuft dendrites during PF formation under *in vivo* conditions. However, the variable magnitude and timing of their recruitment raised the intriguing possibility that PF-forming plateau potentials do not typically originate within the distal tuft compartment as previously thought^40, 57, 61^.

**Figure 4.**
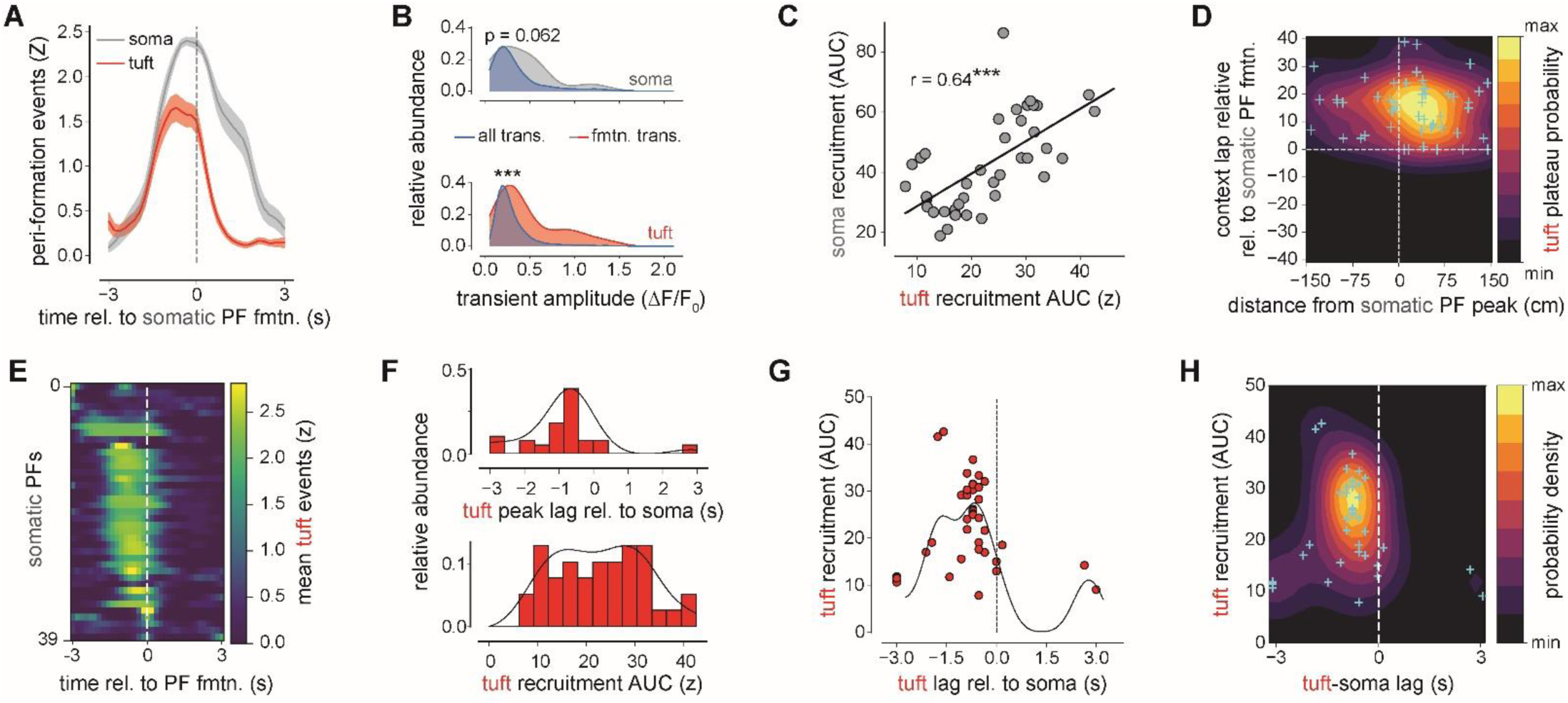
Variable timing and magnitude of distal tuft dendritic activity during spontaneous place field formation. (**A**) Somatic (grey) and tuft (red) peri-event time histograms (PETHs) triggered on somatic PF formation events. Shaded regions indicate SEM. Dashed line indicates estimated moment of PF formation (see supplementary materials and methods). (**B**) Gaussian kernel density estimates (KDEs) showing distributions of all *Ca^2+^*transients (blue) compared to those occurring during PF formation for soma (top, grey) and tuft dendrites (bottom, red). (**C**) Scatterplot showing area under the curve (AUC) of peri-formation PETHs (as in A) for soma (y-axis) and tuft dendrites (x-axis). Black line represents ordinary least squares regression fit. Pearson correlation coefficient is shown. (**D**) Tuft plateau probability as a joint function of lap relative to PF formation (y-axis) and within-lap distance from somatic PF peak (x-axis). Cyan crosses indicate real data points and heatmap depicts joint Gaussian KDE. (**E**) Heatmap of tuft PETHs triggered on somatic PF formation events, i.e. data underlying red trace in (A). Each row represents the mean z-scored *ΔF/F_0_* of all imaged tuft dendrites for a given PF formation event in a single CA1 PN. (**F**) Histograms and Gaussian KDEs of tuft timing relative to soma (top) and tuft AUC (bottom) during PF formation. (**G**) Scatterplot showing tuft peri-formation AUC (y-axis) and tuft-soma lag (x-axis). Black line represents Gaussian KDE fit to histogram values (not shown) after binning AUC according to lag. (**H**) Joint probability distribution of peri-formation tuft AUC and lag relative to soma. Cyan crosses indicate true data points used to fit the heatmapped joint Gaussian KDE. \*\*\**P* < 0.001. See table S1 for sample sizes and table S2 for additional statistical details.

To specifically probe the role of tuft-originating plateau potentials, as opposed to ordinary *Ca^2+^* transients, in PF formation, we assessed their prevalence with respect to lap number and spatial location. This analysis revealed that very few PF formation events were accompanied by tuft plateaus (2 from 39 formation events and 44 plateau potentials within the same virtual context). Instead, tuft plateaus tended to occur after somatic PF formation and slightly forward in space relative to somatic PF peaks (µ = 20.01 ± 10.93 cm, σ = 74.92, *N* = 48 plateaus from 46 tuft dendrites) (Figure 4D). The occurrence of tuft plateaus after PF-associated somatic events may be explained by a variety of factors including ongoing local excitatory synaptic input, local tuft plasticity induced during somatic PF formation, and/or centrifugally-propagating somatic activity during PF traversals. We note that the forward-shifted nature of post-formation tuft plateaus relative to somatic PFs is reminiscent of the ‘peak shift’ characteristic of the BTSP mechanism (Figure 3F) ^39^, whereby somatic PFs emerge at locations the animal occupied 1-3 s prior to plateau initiation, and consistent with a previously suggested role for plateau potentials in boosting the gain of extant somatic PFs^57^.

Given the surprising lack of association between tuft plateaus and PF formation, we more closely examined peri-formation tuft activity profiles across PF formation events. We observed considerable variability not only in the magnitude of peri-formation tuft activation (µ = 22.96 ± 1.47 area under the curve, σ = 9.064, *N* = 39 PF formation events), but its timing relative to the moment of peak peri-formation somatic activity (med. = - 0.70 sec, IQR = 0.53, *N* = 39 PF formation events) (Figure 4, E & F). Remarkably, the distribution of tuft-soma timing lags was identical in both shape and time course to the currently unexplained asymmetric plasticity kernel characteristic of BTSP^39, 42, 43^. The magnitude and timing of peri-formation tuft activity clustered together such that distal tuft dendrites tended to display the strongest activation within 1.5 seconds preceding somatic PF formation (Figure 4, G & H). These data are consistent with prior *in vitro* experiments indicating that properly timed cortical input is critical to drive plateau potentials in CA1 PN distal tuft dendrites^57, 61^. However, the diffuseness of this clustering and the lack of tuft-associated plateau potentials during somatic PF formation show that CA1 PNs can accommodate a wide range of tuft activity patterns in generating new PFs.

In summary, while CA1 PN distal tuft dendrites appear to play a prominent role in PF formation, they do not appear to do so via locally generated plateau potentials. From these results, we conclude that (1) PF- forming plateau potentials originate elsewhere in the CA1 PN dendritic arbor (e.g. the apical trunk within *stratum radiatum* or near the nexus), (2) plateaus do not regeneratively propagate to drive global *Ca^2+^*influx throughout CA1 PN dendritic arbor, and (3) given that tuft-associated plateaus nonetheless drive robust somatic responses (Figure 3, L & M), they primarily serve to boost the expression of existing somatic PFs.

### The timing and magnitude of distal tuft recruitment jointly determine properties of new place fields

The prominent variability in tuft activity during PF formation prompted us to hypothesize that it may shape aspects of new PFs. We sought to (1) test our hypothesis analytically by devising a model to predict PF properties based on peri-formation tuft recruitment and (2) interrogate this model to understand the logic by which tuft dendrites might shape PF properties. Visually inspecting the timing and magnitude of peri-formation tuft recruitment relative to various properties of PF expression and formation revealed several apparent trends with varying degrees of nonlinearity (Figure S9). These trends presented a challenge as they complicated the use of easy-to-interpret linear regression models. Applying a nonlinear kernel to a linear model was inappropriate as we could not reasonably select and apply one polynomial order to both of our tuft features of interest (Figure S9). We opted for an agnostic approach, transforming both tuft recruitment features into 3^rd^ order polynomial feature space and allowing a regularized linear model to decide the relative importance of polynomial features corresponding to *N^th^* order coefficients (Figure 5A).

**Figure 5.**
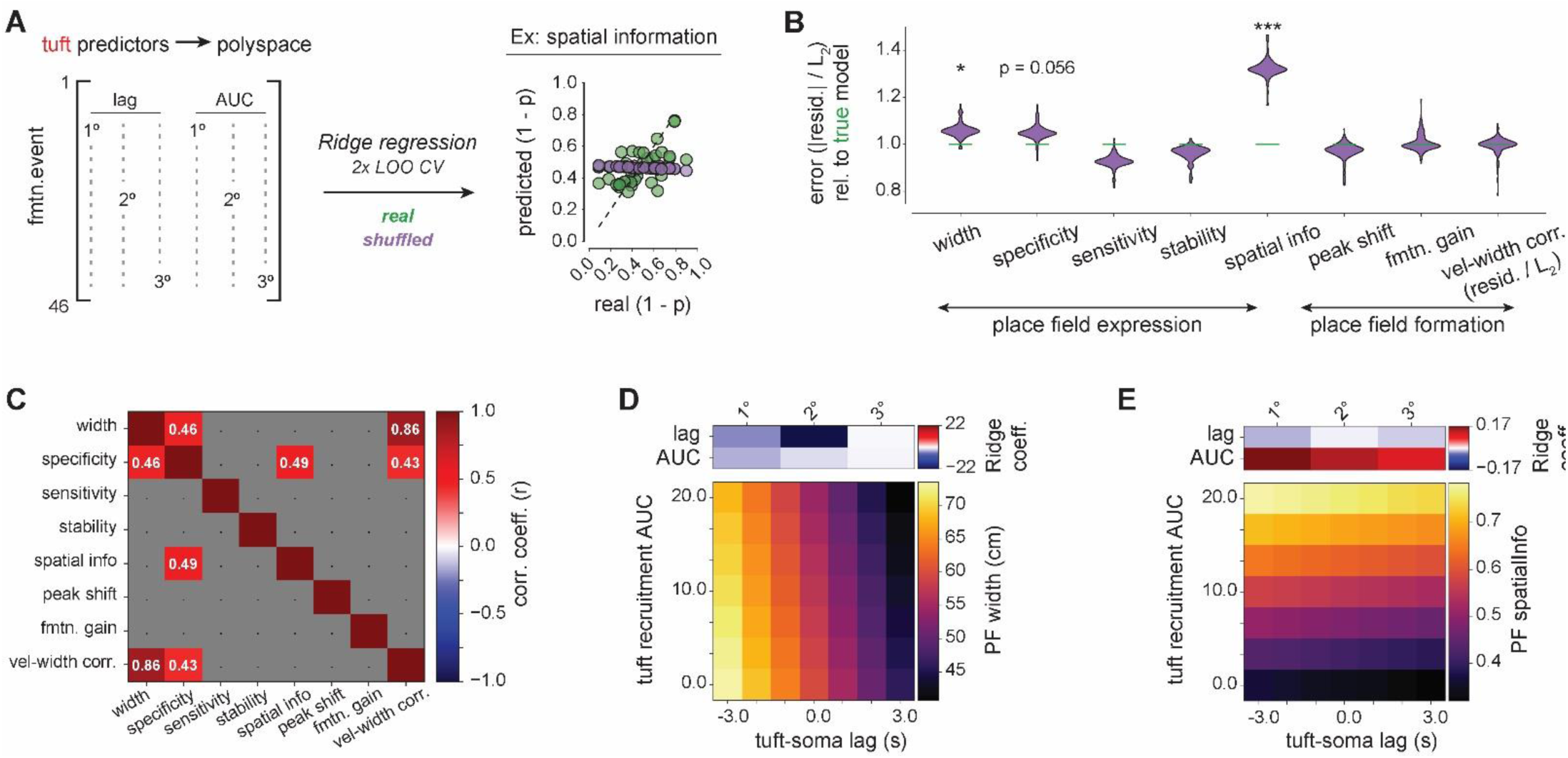
Peri-formation tuft recruitment closely predicts properties of place field expression and formation. (**A**) Schematic describing approach to predict somatic PF properties using features of peri-formation tuft recruitment. Tuft-soma lag and area under the curve (AUC) were transformed into polynomial feature space to train a double-cross-validated Ridge regression model (see supplementary materials and methods). Example result for PF width shown at right with predicted values using real (green) and shuffled (purple) data. Dashed line represents equality. (**B**) Tuft-based model prediction error for properties related to the expression and formation of new somatic PFs. Prediction errors from 1,000 models trained on shuffled data (purple) were normalized and compared to those from real data (green). Violin bodies span full data range with lines at medians and error bars spanning first to third quartiles. (**C**) Correlation matrix showing Pearson coefficients between target variables in (B). Grey cells indicate non-significant coefficients (*P* > 0.05). (**D, E**) Model predictions that significantly outperformed those of models trained on shuffled data, including PF width (D) and spatial information content (E). Top heatmaps in each panel show Ridge model coefficients by tuft feature (rows) and polynomial order (columns). Bottom heatmaps show model predictions across interpolated feature space from Ridge regression models trained on real data. \**P* < 0.05, \*\*\**P* < 0.001. See table S1 for sample sizes and table S2 for additional statistical details.

The magnitude and timing of tuft recruitment during PF formation predicted the resultant width of new PFs and, in particular, their spatial information content (Figure 5, A & B). Peri-formation tuft dynamics did not predict hallmarks of the BTSP mechanism that have been previously established and independently reproduced^20, 39, 43, 82^, including in this study (Figure 3, E & F). Among these hallmarks is a linear relationship between an animal’s running speed during PF formation and the width of the resulting PF (Figure 3E), presumably due to velocity-modulated CA3 inputs recruited a larger proportion of CA1 PN synapses within the seconds-long BTSP plasticity window. Notably, peri-formation tuft activity represents a novel determinant of PF width; neither the magnitude (*R* = -0.18, *P* = 0.26) nor the timing (*R* = 0.28, *P* = 0.083) covaried with peri-formation animal velocity. Finally, despite varying degrees of multi-collinearity among PF properties (Figure 5C), regression weights indicated that tuft recruitment predicts these properties according to unique rules. PF width was most strongly predicted by the timing of peri-formation tuft recruitment (Figure 5D) whereas PF spatial information was almost entirely predicted by the magnitude of tuft activation (Figure 5E). Exact regression weights are shown in table S3. In summary, this analysis identifies the timing and magnitude of tuft activity, surrounding the moment of somatic PF formation, as previously unappreciated determinants of the hippocampal spatial code.

### Local distal tuft spatial tuning is robust and shifted in space relative to somatic tuning

Distal tuft dendrites shape the formation of new PFs while firing plateau potentials during subsequent PF traversals. However, it remains unclear (1) whether tuft dendrites themselves display local PFs and (2) how dendritic PFs hundreds of microns from the soma might relate to somatic PFs. For instance, CA1 PN radial oblique dendrites, located below the distal tuft region within *stratum radiatum*, display PFs with no apparent relation to their somatic PF^20^. CA1 PN distal tuft dendrites undergo local, long-term potentiation *in vitro^83^* and receive spatially informative cortical inputs^40, 47, 48, 55, 56, 58, 84^ via clustered synaptic connections rich in plasticity-promoting *N*-methyl, *D-*asparate receptors (NMDARs) ^72^. We therefore hypothesized that distal tuft dendrites express local PFs. We further hypothesized that, if tuft dendrites do express local PFs, then they may be formed in tandem with somatic PFs and would therefore be tuned to similar locations.

To examine distal tuft spatial tuning relative to somatic tuning, we analyzed data from single-context experiments (Figure 1, C & D) which provided sufficient numbers of laps (µ = 107.25 ± 12.94, σ = 34.24, *N* = 8 imaging sessions from 7 animals) for robust spatial tuning curve (TC) calculations in all cells (Figure S10). Consistent with our hypothesis, 90 ± 4.8% (σ = 13%, *N* = 46) of all imaged tuft dendrites displayed PFs (Figure 6, A & B) with similar properties to somatic PFs (Figure S11). Tuft dendritic PFs were systematically shifted backward in space relative to somatic PFs by 21.43 ± 11.77 cm (σ = 68.60 cm, *N* = 35 tuft-soma PF pairs) (Figure 6, A & C) with a heavy distribution tail at lower absolute distances (Figure 6D). We note that this backward shift is consistent with the tendency for distal tuft dendrites to fire before their soma during somatic PF formation (Figure 4F). Taken together, these result suggests that CA1 PN distal tuft dendrites form local PFs in the process of driving somatic PF formation and their tendency to fire before the soma systematically shifts their spatial tuning preferences backward in space.

**Figure 6.**
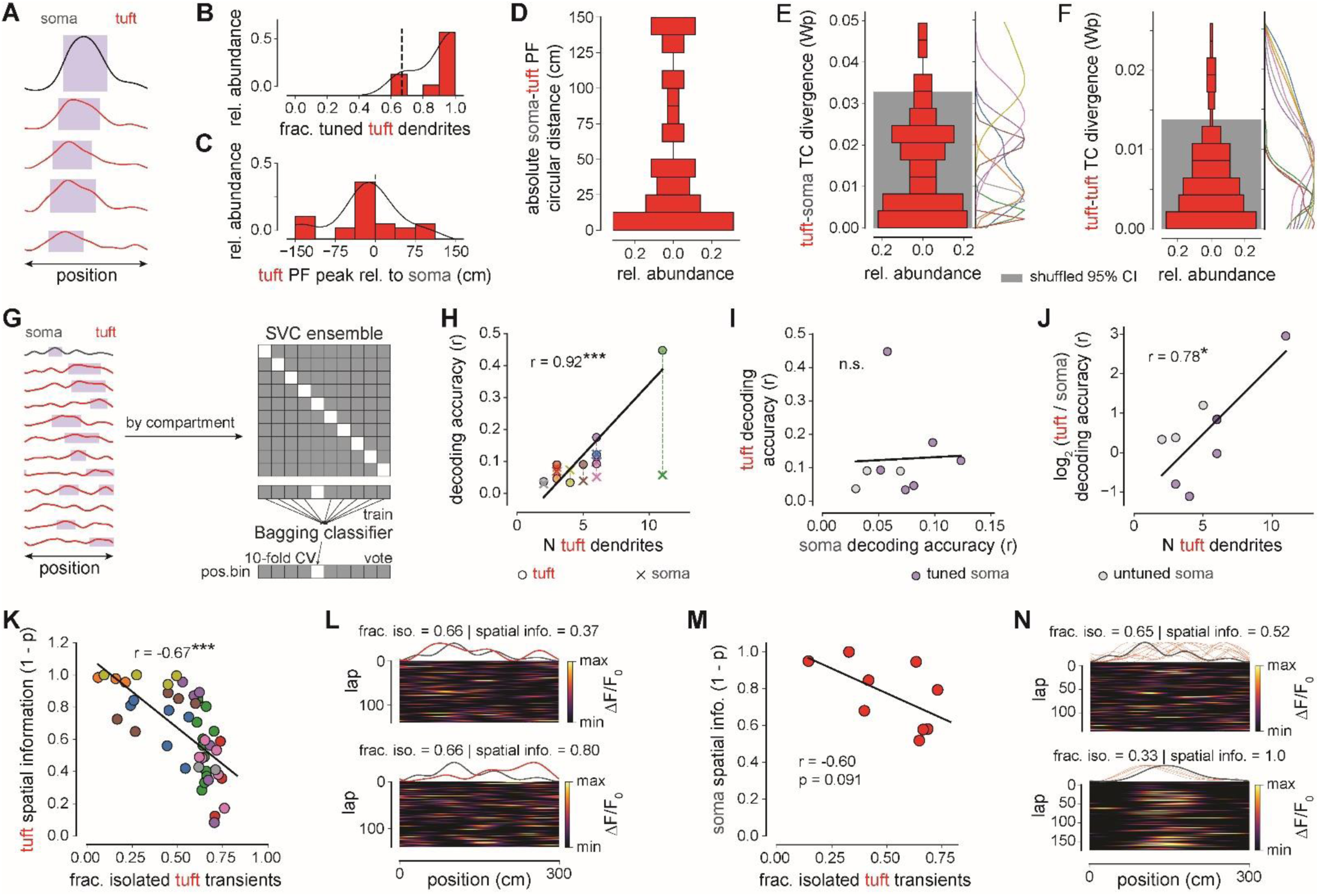
Local spatial tuning in distal tuft dendrites of CA1 PNs. (**A**) Somatic (black) and distal tuft dendritic (red) TCs. Violet shaded regions indicate PFs. (**B**) Histogram and Gaussian kernel density estimate (KDE) of fraction of tuft dendrites per cell expressing PFs. Dashed line indicates fraction of soma expressing PFs. (**C**) Histogram and Gaussian KDE of tuft dendritic PF peak location relative to that of soma. Dashed line indicates equivalent PF peak locations. (**D**) Vertical histogram of absolute circular distances separating soma and tuft PFs. 150 cm is the maximum possible distance along a 3-m virtual track. (**E-F**) Vertical histograms showing divergence (Wasserstein distance, *W_p_*, see supplementary materials and methods) between all connected soma-tuft (E, *N* = 46 pairs from 9 cells) or tuft-tuft (F, *N* = 160 pairs from 9 cells) pairs. Shaded regions indicate 95% confidence interval (CI) generated from TCs calculated on shuffled data. By-cell Gaussian KDEs at right reflect within-cell variability. (**G**) Diagram of support vector classifier (SVC) ensemble-based approach to decode animal position from soma or tuft TCs (see supplementary materials and methods). (**H**) Decoding accuracy plotted against the number of tuft dendrites used for training. Circles represent tufts, Xs separated by dashed lines indicate connected soma, colors indicate cell identity. Black line represents ordinary least squares fit. (**I**) Tuft decoding accuracy plotted against soma decoding accuracy. Violet symbols indicate cell with somatic PF; grey symbols indicate no PF. (**J**) Tuft decoding outperformance relative to connected soma (log_2_ ratio) plotted against number of tuft dendrites used for training. (**K**) Tuft spatial information score as a function of the fraction of its *Ca^2+^* transients that were isolated. Colors correspond to cell identity. (**L**) Example spatial tuning heatmaps showing tuft *ΔF/F_0_* as a function of lap number (y-axis) and animal position (x-axis). Tuft (red) and soma (black) TCs are plotted above. Top: Example consistent with trend shown in (K). Bottom: Counterexample to trend shown in (K). (**M-N**) Soma spatial information score, as shown in K and L. \**P* < 0.05, \*\*\**P* < 0.001. See table S1 for sample sizes and table S2 for additional statistical details.

Since soma and dendrites lacking statistically significant PFs nonetheless displayed subthreshold tuning preferences with apparent cross-compartment relationships (Figure S10), we next compared spatial TCs using pairwise Wasserstein distances. In this setting, Wasserstein distance describes the amount of ‘work’ required to transform one spatial TC into another (e.g. lower values indicate that two TCs are more similar to each other). Consistent with our hypothesis, somatic and distal tuft spatial tuning preferences were generally similar with few tuft TCs significantly differing from those of their soma (Figure 6E). The same was true between individual tuft dendrites of a given cell (Figure 6F), although we noted several exceptions to both of these trends (Figure S10; see also by-cell distributions in Figure 6, E & F). Taken together, these data show that distal tuft dendrites tend to display PFs back-shifted in space relative to somatic PFs, which is accounted for by their activity during somatic PF formation, but they nonetheless have the capacity to express fully autonomous spatial tuning preferences.

Given the non-negligible variability of tuft spatial tuning preferences across and within CA1 PNs, we next asked whether tuft dendrites locally encode spatial information beyond that which might be inherited from backpropagating somatic APs. To this end, we trained a support vector classifier ensemble to predict each animal’s position along the 3-m virtual track based on either tuft or somatic TCs from a single CA1 PN (Figure 6G). To be clear, this model was destined to perform poorly given that it was trained on either 1 soma or 2-11 tuft dendrites (see table S1). Rather than accurately predict animal position, our goal was to assess per-tuft spatial information content relative to the soma. This analysis revealed that tuft dendrites linearly sum to represent an animal’s environment such that the number of dendrites used for model training predicts the extent to which the distal tuft compartment outperformed the soma in decoding animal position (Figure 6, H-J). These results demonstrate that not only are distal tuft dendrites spatially tuned, but their diversity in spatial tuning preferences renders them collectively more spatially informative than the soma.

Finally, although tuft dendrites autonomously encode animal position, this does not appear to be their primary function. To the contrary, a tuft dendrite’s overall fraction of isolated *Ca^2+^* transients negatively predicted its spatial information content (Figure 6K, see Figure 6L for example and counterexample). A similar but non-significant trend was observed at the soma (Figure 6, M & N). These data indicate that, despite the high prevalence of PFs in tuft dendrites (Figure 6B), and their cumulatively robust spatial information content (Figure 6, H-I), isolated tuft *Ca^2+^* transients primarily encode non-spatial information. To better understand the information content of isolated tuft transients, we attempted to relate isolated tuft activity to a collection of locomotor and reward-related behavioral features using a cross-correlation-based approach (see supplementary materials and methods). Briefly, tuft *ΔF/F_0_* signals not corresponding to isolated *Ca^2+^* transients were masked with zeros. As a control, the same was done for somatic signals not corresponding to a somatic *Ca^2+^* transient. Most soma and tuft dendrites were significantly correlated to all features that we assessed: velocity, acceleration, deceleration, rewarded licks, and unrewarded licks (Figure S12, A & B) with variable time lags (Figure S12C). Isolated tuft events were more behaviorally informative than somatic events on the whole, with no one behavioral feature standing out (Figure S12D). In summary, distal tuft dendrites carry out diverse local operations hundreds of microns from their soma. They possess robust, local PFs that tend to be locally similar to each other, and globally back-shifted relative to the somatic PF, while still possessing potentially consequential diversity in their tuning preferences. Finally, while isolated tuft activity can clearly contribute to local spatial tuning, it primarily encodes non-spatial behavioral features.

## DISCUSSION

By simultaneously monitoring somatic and distal tuft dendritic *Ca^2+^* dynamics *in vivo* during two virtual spatial navigation tasks, we have uncovered several previously unappreciated aspects of CA1 PN distal tuft dendritic function. Given the lack of prior knowledge regarding distal tuft dynamics in CA1 PNs *in vivo*, our study is necessarily broad and descriptive. Here we wish to provide additional context to synthesize our findings and to reconcile key observations with the current model for BTSP-driven PF formation.

We find that distal tuft dendrites are strongly compartmentalized from their soma in CA1 PNs and favor centripetal propagation of dendrite-originating events. While strong compartmentalization was largely expected based on past *in vitro* and computational work^60, 70-73, 77, 85^, we were surprised to find that compartmentalization broke down during immobility. CA1 PN soma are more active during locomotion and somatic depolarization results in better soma-tuft coupling *in vitro* due to A-type *K^+^* channel inactivation^73^. At the circuit level, velocity-modulated EC neurons transmit increased excitatory signals to tuft dendrites during running states^54^. Indeed, this is consistent with our observation that tuft dendrites are far more active during running than immobility. Furthermore, EC-driven tuft activation should be boosted by clustered^86^, NMDAR-rich^72^ EC-tuft synapses and depolarization-dependent inactivation of *I_h_* currents^70, 87-89^. These studies all point to increased soma-tuft coupling during locomotion.

However, it is also the case that somatic AP backpropagation favors single spikes rather than bursts^90^. Inhibition undoubtedly plays a major role in soma-tuft cross talk as well, via either local disinhibition by long-range interneuron-targeting EC inhibitory projections^91^, depolarizing *I_h_* tail currents^75, 76, 92^ following the release of velocity-modulated ^93^ dendritic inhibition^94-99^, or some combination of both. Therefore, a mixture of inhibitory and cell-intrinsic mechanisms may dynamically modulate distal dendritic excitability as well as its impact on somatic AP firing. We speculate that one function of such gating may be to sparsen the CA1 population code by limiting the rate of PF formation: unchecked plasticity might otherwise bias the hippocampal cognitive map^32, 33^ to disproportionally represent regions first encountered rather than tiling the entirety of an animal’s environment.

Based on our unsupervised plateau detection strategy, which is robust by all measures, tuft dendrites do not form somatic PFs via locally-generated plateau potentials as reasonably postulated based on a substantial amount of converging albeit indirect evidence^39-42, 57, 61^. To be clear, we do not argue against a role for dendritic plateau potentials in somatic PF formation. ADPs are a somatic vestige of dendritic plateau potentials and their presence during PF formation, even in the absence of experimental perturbation^57^, indirectly indicates that they occur during at least a substantial portion of PF formation events. Instead, we suggest that properly timed input converging onto both distal tuft and radial oblique dendrites leads to plateau potential initiation elsewhere in the apical arbor; likely in the apical trunk if not near the nexus. One implication of this revised model is that plateau potentials may not uniformly depolarize the CA1 PN dendritic arbor such that all dendritic spines receiving presynaptic input within a second-long window around plateau onset will undergo plasticity^39^. Intuitively, this is not a problem for the BTSP mechanism: one CA1 PN possesses thousands of dendritic spines and depolarizing a plurality of them may well be sufficient to drive robust PF formation.

Perhaps most intriguingly, we found that CA1 PN distal tuft dendrites exert analog, nonlinear control over the expression of new somatic PFs via the timing and magnitude of their peri-formation recruitment. Given recent evidence that BTSP comprises elements of long-term potentiation (LTP) as well as depression (LTD) ^42, 43^, one attractive possibility is that these features of peri-formation tuft recruitment may influence the balance between LTP and LTD to determine PF fidelity^100^. We also noted that the distribution of tuft timing lags relative to their soma strikingly resembled the asymmetric, seconds-long temporal association window, or ‘plasticity kernel’^39, 42, 43^, that renders BTSP particularly relevant to the timescales of action-outcome relationships encountered in the real world. Efforts are already underway to identify its molecular underpinnings^44-46^. Here we propose that the BTSP plasticity kernel may be specifically tuned to accommodate variable tuft timing. Alternatively, since this kernel is an estimate based on many PF formation events and CA1 PNs, it may simply reflect the temporal variability of tuft recruitment during PF formation. More broadly, we view our results to strengthen the role of distal tuft dendrites in BTSP-mediated PF formation. By virtue of its variable peri-formation recruitment, this distal neuronal compartment may effectively set the gain, or proverbial importance, of new feature-selective receptive fields based on the nature of concomitant, top-down cortical signals.

Finally, tuft activity during somatic PF formation appears to drive local tuft plasticity that subsequently helps to maintain the new somatic PF. Tuft dendrites express local PFs that are back-shifted relative to their somatic PF. The distance of this backward shift is accounted for by the timing differential between tuft and somatic activation during somatic PF formation, indicating that tuft recruitment produces tuft PFs in parallel via local plasticity. As a result, tuft dendrites will be reactivated to a greater extent during subsequent, post-formation traversals of the CA1 PN’s somatic PF. We therefore propose that concomitantly-formed tuft PFs effectively increase the probability of tuft-associated plateau potentials during somatic PF traversals. Given that we found tuft plateaus to strongly drive somatic activity, this would appear to be an elegant mechanism by which tuft dendrites maintain newly formed somatic PFs. In conclusion, we find that distal tuft dendrites constitute an impressively multifunctional neuronal compartment capable of dynamically modulating somatic activity, representing numerous experiential features, and lastingly determining the relative import of new hippocampal receptive fields.

## ACKNOWLEDGMENTS

The authors thank H. Bito for generously providing an XCaMP-R-expressing plasmid DNA construct, J. Bowler and J. Priestley for assistance with virtual navigation, C. Gillon and S. Terada for input on advanced statistical approaches, T. Geiller for productive discussion regarding local CA1 inhibition, and U. Bayer, T. Geiller, M. Szoboszlay, J. Bowler, and K. Gonzalez for invaluable feedback on the initial manuscript. The authors acknowledge support from the National Institutes of Health: K99NS127815 (JKO), R35NS127232 (FP), R01MH124047 (AL), R01MH124867 (AL), R01NS121106 (AL), U01NS115530 (AL), R01NS133381 (AL), R01NS131728 (AL), RF1AG080818 (AL).

## AUTHOR CONTRIBUTIONS

Conceptualization: JKO, FP, AL

Methodology: JKO, SJW, MDS, FP, AL

Investigation: JKO

Software: JKO, SJW

Formal analysis: JKO

Validation: JKO

Visualization: JKO

Data curation: JKO

Funding acquisition: JKO, FP, AL

Resources: FP, AL

Project administration: JKO, FP, AL

Supervision: JKO, FP, AL

Writing – original draft: JKO

Writing – review & editing: JKO, SJW, MDS, FP, AL

## DECLARATION OF INTERESTS

Authors declare that they have no competing interests.

## SUPPLEMENTAL INFORMATION

### Methods

#### Animals

Experiments were conducted in agreement with NIH guidelines and approval by Columbia University’s Institutional Animal Care and Use Committee. Animal welfare was supervised by designated veterinary staff. Columbia University animal facilities comply with all applicable standards of care including cage conditions, space per animal, temperature, light, humidity, nutrition, and hydration. Male and female C57Bl/6 mice were at least 42 days old at time of surgery and no older than 4 months by the conclusion of imaging sessions. Authors are not aware of an influence of sex on the measures of interest to this study.

#### Hippocampal window implants

All procedures were carried out under isoflurane anesthesia on a circulating water heating pad to maintain body temperature. Meloxicam was administered subcutaneously for long-term analegesia and bupivacaine was administered at the incision cite for local, short-term analgesia. Modified imaging windows with a silicon-protected, laser-etched rectangular opening for pipette access were implanted as previously described^20^. Mice received postoperative subcutaneous injections of sterile saline and recovered in home cages with external heat until they recovered sternal recumbency. Mice then received regular postoperative health checks for three days with electroporations carried out no less than seven days post-surgery.

#### Single-cell electroporation

pCAG-XCaMP-R^63^ (gift from Haruhiko Bito, University of Tokyo) was transformed using Stellar Competent Cells (Takara Bio.), grown in antibiotic-treated LB broth at 37° C, and purified using an endotoxin-free midiprep kit (NucleoBond Xtra Midi EF, Macherey-Nagel). Purified plasmid DNA was diluted to a final concentration of 135-150 ng/µL in a solution containing (in mM): 155 K-gluconate, 10 KCl, 10 HEPES, 4 KOH, 0.33 Alexa Fluor 488. Electroporation solutions were calibrated to 7.3 pH and 316 mOsm prior to addition of plasmid DNA and Alexa Fluor 488. Solutions were then successively filtered using 0.45 and 0.20 µm cellulose acetate syringe filters (Invitrogen, 171-0020 & 171-0045). CA1 PNs were then electroporated as previously described^20^. Briefly, anesthetized mice were head-fixed under a 2-photon microscope and a long-tapered borosilicate glass pipette containing electroporation solution was inserted through a rectangular opening in the imaging window that was protected by a thin, penetrable silicon layer. Using a standard electrophysiology amplifier and digitizer to monitor tip resistance, along with 2-photon excitation of Alexa Fluor 488 in the electroporation solution, the tip was guided to CA1 *stratum pyramidale* under continuous positive pressure. When the tip approached a CA1 PN, as judged by deflection in tip resistance, a high-frequency train of negative voltage pulses were delivered to temporarily permeabilize the neuron’s plasma membrane and thus allow plasmid DNA entry. Electroporated neurons were allowed no less than 2 days to express sufficient XCaMP-R protein for 2-photon *Ca^2+^*imaging experiments.

#### Virtual navigation

Water-restricted (≥ 90% baseline body weight), head-fixed mice navigated 3-meter virtual contexts for water reward using a previously described apparatus that updates an array of LCD monitors based on the rotation of a running wheel^56, 67, 82^. During initial training, water was non-operantly delivered at 10 locations spanning the virtual context to establish an association between locomotion and reward delivery. Training ended once mice learned to run for a single reward zone per lap. Running frames were defined as any imaging frame during which the mouse ran at a velocity of ≥ 1 cm / s. LCD screens were blanked during 0.5 – 2.0 s intertrial intervals separating each lap. Data acquired during intertrial intervals were excluded from all analyses.

##### Single-context experiments

In single-context experiments corresponding to Figs. 1, 2, & 6, mice navigated familiar contexts for one pseudorandomly located water reward per lap for 30 minutes. When a mouse entered the reward zone for a given lap, an initial drop of water was non-operantly dispensed via a metal port that detects licking events via a capacitive sensor. Additional licks within the reward zone resulted in operant water delivery for a period of 2 seconds under a fixed ratio 2 reinforcement schedule.

##### Multi-context experiments

Experiments consisting of multiple novel contexts (Figs. 3-5) were used to boost the probability of spontaneous PF formation as novel context exposure is known to drive increased PF formation in CA1 at the population level^80-82^. Mice navigated a series of four novel, 3-meter virtual contexts. The first of these contexts was reused as the familiar environment in the above-described single-context experiments. Mice were ‘teleported’ to the next context after 12 or 15 laps. After mice visited all four novel contexts, they revisited the same contexts in an infinite loop for the remainder of the 30-minute imaging session. In this manner, novel and familiar contexts were internally controlled by context ID when assessing the influence of novelty on subcellular *Ca^2+^*signals.

To encourage the use of visual scenes for navigation, and thereby further boost the probability of spontaneous PF formation, reward zones were positioned at unique, fixed locations in each context for 6/7 mice (6/8 imaging sessions and 7/9 imaged CA1 PNs). Mice underwent additional prior training under the same conditions but with a separate set of novel environments which were subsequently discarded. Narrow (7.5 cm) reward zones required mice to learn reward zone locations and slow down upon approach so as to avoid running through them and forgoing water reward. Mice rapidly learned reward zone locations, as evidenced by anticipatory licking and deceleration (Figure S3). The remaining imaging sessions used pseudorandomly located rewards as described above for single-context experiments and nonetheless resulted in robust spontaneous PF formation (4 events in each CA1 PN). Therefore, mice ostensibly attended to visual scenes regardless of the navigational demands imposed.

#### 2-photon *Ca^2+^* imaging

Individual CA1 PNs were imaged across dendritic and somatic focal planes as previously described^20^ except for additional specifics noted here. XCaMP-R was excited using a Fidelity-2 1070 nm 2-photon fiber laser (Coherent Inc.). Photons were routed by a 575 nm dichroic mirror (575dcxr, Chroma Technology Corp.) through an emission filter (HQ607/45m-2p, Chroma Technology Corp.) to a GaAsP photomultiplier tube (7422P-40, Hamamatsu Photonics K.K.) dedicated for red photon collection. Images were acquired at 512 x 512 pixels with fields of view ranging from 221.25 to 334.12 microns in width. Resolution ranged from 0.43 to 0.65 microns per pixel. A piezoelectric device (Bruker Corp.) was used to rapidly toggle a 40X NIR water immersion objective lens (0.8 NA 3.5 mm WD, Nikon USA) between *stratum pyramidale*, i.e. the somatic imaging plane, and *stratum lacunosum moleculare*, i.e. the distal tuft dendritic imaging plane. The very long jumps between focal planes cause inertial vibrations in the objective lens upon stoppage, which then leads to undesirable imaging artifacts. Introducing a 65 ms ‘wait time’ between toggling the axial position of the objective lens and acquiring imaging data consistently circumvented these artifacts while permitting sufficient cross-plane frame rate (∼6 Hz) to acquire high-quality XCaMP-R signals (Figs. 1D & 3H).

The same CA1 PNs were imaged across both behavioral paradigms. To image the same focal plane of dendrites in each experiment, time-averaged fields of view (FoVs) from 30-60 s test recordings were compared to motion-corrected, time-averaged FoVs from the prior experiment and piezoelectric device parameters were iteratively tuned until these images were indistinguishable. Imaging data were motion-corrected using a recently described iterative, template-matching approach designed for relatively dim signals^20^. Regions of interest (ROIs) corresponding to soma and distal tuft dendrites were hand-drawn over motion-corrected, time-averaged FoVs.

Regions of interest (ROIs) were manually drawn and annotated to reflect their position in the CA1 PN dendritic arbor, as determined from z-stack images of each CA1 PN acquired at 1-micron axial resolution under general anesthesia. Each dendritic ROI corresponded to the entire cross section of a unique dendritic branch that was present in the field of view (Figure 1B). Multiple ROIs were never drawn for the same dendritic branch. Dendritic anatomical features were not a major focus of the present study as path lengths separating distal tuft dendrites from their soma are uniformly long and branch order was not found to predict measures of interest. Nonetheless, one CA1 PN was digitally reconstructed for illustrative purposes. To this end, a custom Python script was used to denoise 3D images and correct for depth-related changes in signal intensity that were not corrected by laser power compensation (Figure 1E). The processed 3D image was then automatically traced, filled, and segmented (Figure 1F) using the SNT FIJI plugin^101^ and a custom Jython script.

#### Signal processing

##### ΔF/F_0_ calculation

To calculate *ΔF/F_0_* traces from raw *Ca^2+^* signals, raw signals were first assessed for frames in which motion correction failed. An initial baseline signal was calculated as the rolling maximum of rolling minimums (max-min) using a Gaussian smoothing kernel (σ = 10 s). Frames in which an ROI’s signal dropped below baseline by at least 1 standard deviation of the above-baseline signal were masked. Masked signals were then denoised using a Python adaptation of the MATLAB (The MathWorks, Inc.) wavelet denoising implementation (wden). Masked data points were sliced out of signals prior to denoising and then reinserted afterwards to avoid corrupting wavelet transforms. A final baseline calculation was then run on denoised data using a σ = 1s Gaussian smoothing kernel. Final *ΔF/F_0_* was calculated as (raw – baseline) / baseline and smoothed using a σ = 0.5 s Gaussian smoothing kernel while skipping any NaN values at the beginning or end of the trace. Fluorescence dynamics were z-scored and reported as such when used for machine learning approaches; in all other cases, traces are displayed as true *ΔF/F_0_*.

##### Event deconvolution

‘Spike’ deconvolution was used to generate binary event trains for spatial tuning analysis. Deconvolved events were not considered or treated as true action potentials or dendrites spikes; rather, binary event trains simply allow for more straightforward identification of activity that is spatially tuned above chance levels. Deconvolved events were calculated from *ΔF/F_0_* traces using the OASIS deconvolution software package^102^ as previously described^20^ with a 2-MAD threshold, 400 ms decay constant, and max-min baseline with a σ = 5 frames Gaussian smooth kernel.

##### Ca^2+^ transient detection

*Ca^2+^* transients were identified using a template matching strategy. The traditional use of an amplitude cutoff based on signal variance typically misses small events because the cutoff must be set well above noise to avoid false positive detections. In contrast, template matching robustly identifies events regardless of amplitude while correctly rejecting events of similar amplitude if their waveforms do not resemble the template, i.e. expected *Ca^2+^*transient waveform. To generate an initial *Ca^2+^* transient template waveform, transients were identified using a simple standard deviation cutoff of 1.5 σ for transient onset and 0.5 σ for transient offset, as well as a minimum duration of 0.25 s. A relatively stringent 1.5 σ minimized false positive detections to generate an accurate template. Peri-event time histograms (PETHs) were then calculated for these initial transients with a window spanning 1 s before and 3s after transient peak. PETHs were pooled and averaged without smoothing to obtain a template waveform. This process was performed separately between cellular compartments (soma and distal tuft dendrites) and datasets (Figs. 1, 2, & 6; Figs. 3-5). *ΔF/F_0_* traces were then continuously correlated to transient template waveforms using a sliding window to generate a like-sized array of correlation coefficients. A peak detection algorithm^103^ was run on the coefficient array with a peak threshold of *R* = 0.85 to identify preliminary events. Finally, preliminary events were bounded using a symmetric threshold of 1 σ, minimum duration of 1 s, maximum duration of 30 s, and lockout period of 0.5 s. As previously described^20^, final traces and transients were obtained after 3 iterative cycles of baseline re-estimation while masking transients identified in the previous round.

##### Plateau potential detection

To identify putative plateau potentials in distal tuft dendrites in an unsupervised manner, all *Ca^2+^* transient PETHs (z-scored, aligned by peak, and offset to zero minimum values) were first factorized using a 2-component non-negative matrix factorization (NMF) model. A *K*=3 K-means clustering model was then fit to the factorization matrix. The cluster with the highest mean peak amplitude was attributed to plateau potentials and the remaining two clusters were merged. Events that were classified as plateaus but did not exceed the 95^th^ percent of maximum amplitudes in the transient (non-plateau) cluster were redesignated as ordinary *Ca^2+^* transients. A plateau potential template waveform was then generated as described above for *Ca^2+^* transient detection using z-scored PETHs of all classified plateau potential events.

Once a plateau potential template *Ca^2+^* waveform was generated, an ensemble of 10 support vector classifiers with *L_2_* regularization parameters of 1.0 was used to differentiate between plateau potentials and *Ca^2+^* transients based on z-scored PETHs. 10 classifier ensembles were trained on random subsets of events (33% for soma, 90% for tuft dendrites) and tested on the remaining held out data. PETHs were resampled as necessary due to minor differences in sampling rates between imaging sessions which arose from variations in piezoelectric device jump size. The classifier ensemble with the highest accuracy (all above 99%, Figure S6D) was saved and linked to a given dataset. With a plateau classifier in hand, template-based *Ca^2+^* transient detection was rerun using both templates. Duplicate events were discarded and overlapping events were merged. The trained support vector classifier ensemble then labeled each event as either a *Ca^2+^* transient or a plateau potential. Finally, an all-or-none condition was imposed such that, for a putative plateau to be included in future analyses, all tuft dendrites of a given cell should be active (at least a *Ca^2+^* transient) and at least 50% of their events should have been classified as plateau potentials. If this condition was met, then each coactive event was considered a plateau potential. If this condition was not met, then each coactive event reverted to a *Ca^2+^* transient. Soma classifications were independent of tuft classifications. Scikit-learn^103^ implementations of NMF, K-means clustering, support vector classification, and bagging classifier were used.

#### Comparing somatic and distal tuft *Ca^2+^* dynamics

##### Cross-compartment propagation of Ca^2+^ transients

Dendritic *Ca^2+^* transients were defined as ‘isolated’ if their bounds did not intersect with those of any *Ca^2+^*transient in their connected soma (e.g., Figure 2B). Conversely, two ROIs were considered coactive for a given event in ROI-A if ROI-B displayed a *Ca^2+^* transient with overlapping bounds. In conditional analyses (Figure 2C), the same logic was applied to calculate what fraction of ROIs from a given cellular compartment displayed *Ca^2+^* transients that overlapped with each of a given ROI’s *Ca^2+^* transients. To determine relative time lags between tuft and soma ROIs (Figure 2D), tuft signals were averaged within-cell and transient detection was rerun on mean signals. Somatic *ΔF/F_0_* was then triggered on the onset of each detected mean tuft event only if the soma fired a *Ca^2+^*transient that overlapped in time with the tuft *Ca^2+^* transient that was used as a trigger. Tuft signals were averaged to guard against that fact that *Ca^2+^* transients in the distal tuft compartment recruited variable fractions of individual tuft dendrites (Figure 2C) since this variability may impact the timing and magnitude of somatic responses.

##### Predicting somatic ΔF/F_0_ from distal tuft dendrites

To predict somatic *ΔF/F_0_* from that of connected distal tuft dendrites (Figure 2E), we trained a 10-fold double-cross-validated Ridge regression model^103^ on 100 random subsamples of *N* dendrites where *N* ranged from 1 to the number of dendrites imaged for a given cell or the maximum *N* that was imaged in at least two CA1 PNs; whichever was least. *ΔF/F_0_* traces were scaled to [0, 1] before model training and testing. The top-level cross-validation step was circular such that non-overlapping chunks of 10% of test data were iteratively predicted to produce one full fit. The mean cross-fold *R^2^* accuracy score was stored for each *N* along with the mean fit from full models which were trained on all imaged dendrites. This approach was identical to that previously used to decode somatic dynamics from those of basal and radial oblique dendrites in CA1 PNs^20^.

#### Spatial tuning analysis

##### Tuning curves and place fields

Spatial tuning curves (TCs), place fields (PFs), and PF properties were calculated as previously described^20^ with the exception that 150 position bins were used with a 7.5-bin Gaussian smoothing kernel. These values were increased proportionally to the 3 m virtual track length compared to a previous 2 m fabric treadmill belt. To calculate spatial tuning features in multi-context experiments, traversals of a given context were concatenated.

##### Comparing spatial tuning profiles

Spatial tuning preferences were compared between ROIs using two metrics: PF circular distance (Figure 6, C & D) and Wasserstein distance (Figure 6, E & F). Circular distance described the minimal distance, in either direction, separating the peaks of two PFs along a 3 m circular track. The maximum distance was therefore 150 cm. When ROIs displayed multiple PFs, circular distances were computed between the most proximal PF pairs. Wasserstein distance was used as a complementary metric that did not require the presence of a statistically significant PF. This metric was deemed valuable as subthreshold spatial tuning features were readily apparent in all imaged somatic and distal tuft ROIs (Figure S10). Compared to standard correlative analyses, Wasserstein distance is resistant to systematic offsets (Figure 6C) in otherwise-similar TCs as it describes the minimal ‘effort’ required to reshape one data array to match another. Since Wasserstein distance values are relative and therefore uninformative without a reference point, 1,000 TCs calculated from shuffled data were used to generate 95% confidence intervals. Shuffled TCs were identical to those used to calculate statistical significance of PFs as they were stored at the end of PF detection.

##### Decoding animal position from spatial tuning curves

Animal position was decoded from TCs of either soma or distal tuft dendrites for each imaged CA1 PN (Figure 6, G-J) using ensembles of 100 support vector classifiers with 10-fold cross-validation and *L_2_* regularization parameters of 1.0. Animal position was indexed to 1 of 150 position bins across a 3 m virtual track for each imaging frame to generate an *N*-frame array of position bin values. TCs were scaled to [0, 1] prior to model training and testing. Cross-validated models were circularly fit to 90% of these data and tested on the held-out 10% to produce full predictions of binned animal position. Each of these top-level folds were repeated 10 times to generate mean fits and *R^2^* accuracy scores. The scikit-learn module^103^ was used to create a pipeline with built-in scaling that contained a bagging classifier of support vector classifiers.

#### Relating *Ca^2+^* signals to non-spatial behaviors

##### Cross-correlation analyses

To assess the correlation between ROI *ΔF/F_0_* and non-spatial behaviors (Figure S12), cross-correlation was used to identify the maximum lagged Pearson correlation coefficient (*R*). *ΔF/F_0_* and behavior traces were scaled to [0, 1] prior to cross-correlation. Correlation coefficients were normalized with the product of vector norms to obtain *R* values on the interval [-1, 1]. Dendritic *ΔF/F_0_* traces were set to zero expect for imaging frames corresponding to isolated *Ca^2+^* transients. Somatic *ΔF/F_0_* traces were set to zero expect for imaging frames corresponding to any somatic *Ca^2+^* transient. Frame-by-frame values for velocity, acceleration and deceleration were used. Acceleration and deceleration arrays contained only positive and zero values. Running was defined as a Boolean array based on a running threshold of 1.0 cm /s. Rewarded and unrewarded licks were defined as Boolean arrays using behavior onset frames. Behavioral data were smoothed with the same 0.5 sec Gaussian kernel used to smooth *ΔF/F_0_* traces. A maximum lag of 5 s was allowed in either direction. This value was determined by inspecting peak lag values, i.e. time lags indexed by maximum *R* values. Time lags were gradually increased from ± 2 s until few ROIs displayed maximum *R* values at the extrema of the allowed temporal window. Frames masked during motion correction were sliced out consistently between behavior and *Ca^2+^* signals such that peak lag calculations were unaffected. The fraction of ROIs with *R* values significantly above chance (Figure S12B) was determined using 1,000 *R* values from shuffled data and a 95% confidence interval.

#### Newly formed place fields

##### Detection

To detect *de novo* PF formation events, spatial tuning was analyzed over concatenated traversals for each context as described above. For each of four unique contexts, putative PFs were calculated as described above but with relaxed confidence intervals. All PFs were considered to be newly-expressed as each context that the animal encountered was novel upon first traversal (Figure 3A). The default alpha of 5% was multiplied by the number of traversals of a given context for a minimum confidence interval of 100 - alpha. This permissive significance threshold was used as a first-pass strategy to account for the fact that PFs may be traversal-specific, form relatively late in the concatenated context, or disappear midway through. In any of such cases, true PF formation events would likely have been missed as the PF would not have expressed across sufficient laps. In this first-pass detection step, the confidence interval was set to 95% and successively stepped down in increments of 10% to 100% - (5% x *N*_traversals_) with any newly-discovered PFs, i.e. PFs not overlapping with any already identified, stored.

For each putative PF, a separate algorithm attempted to identify the precise moment of PF formation. First, the formative lap was identified as the first lap in which activity within PF bounds (‘hits’) exceeded the mean of non-zero hit values. We note that this approach biases our detection strategy toward BTSP-mediated PF formation events which are characterized by a particularly high degree of somatic activity relative to subsequent PF traversals^39, 43, 82^ (‘formation gain’, see Figure 3F). For PF formation events occurring after the first 8 laps of a given context, post-formation PF hits were required to exceed those in pre-formation laps as determined by *t*-test. This test was not run on PFs that formed within the first 8 laps of novel context exposure as insufficient pre-formation laps were available for meaningful statistical comparison.

Within the now-identified PF formation lap, the precise imaging frame of PF formation was defined as the imaging frame in which *ΔF/F_0_* exceed the 99^th^ percentile of in-PF values on the formative lap. Animal position was then indexed to this frame to identify the spatial location of the PF formation event. The forward PF bound was expanded by 10% of the virtual track length (30 cm) for this step to account for the documented backward ‘peak shift’ characteristic of BTSP-mediated PF formation^20, 39, 43, 82^. If a separate PF was found to occur within the same context, the current PF’s hit rate was compared before and after the formation lap of the secondary PF. This step accounted for the possibility that PFs may ‘remap’, i.e. disappear upon the emergence of a new PF. In this case, a ‘stop lap’ was attributed to the first PF and used when calculating PF properties (described below). Otherwise, the stop lap was assigned as the final lap of the concatenated context. If a formative event for a putative PF could not be identified (e.g. pre-formation PF hits were not significantly greater than post-formation PF hits), the PF was discarded.

Putative PFs with identified formation events were then further validated taking into account the laps at which the PF formed and, if applicable, disappeared. Validation was less stringent than for single-context experiments with many laps (Figs. 1, 2, & 6) which exclusively used 95% confidence intervals relative to TCs calculated from shuffled data, but was more rigorous than initial identification of putative PF formation events. For each context, a minimum confidence interval was set to 100% - (5% x *N_events_*) with *N_events_* referring to the number of putative PF formation events detected for a given PN in a given context. Confidence was stepped down from 95% to this minimum in 5% increments with significant PFs that overlapped with the putative PF by stored at each interval. If multiple PFs overlapped with the newly-formed PF of interest, the PF with greatest similarity to the putative PF was stored and the other(s) ignored. Finally, since PF validation typically returned PFs with slightly different bounds than initial putative PFs, formation lap and imaging frame/position were recalculated as described above.

##### Metrics

Several metrics were calculated for each identified PF formation event. Metrics describing PF expression (sensitivity, specificity, width, and spatial information score) were calculated as described above, using only those laps ranging from PF formation to the final PF lap. PF stability (Figure 5; Figure S11) was quantified as the log_2_ ratio of PF sensitivity during the first re-exposure to the context in which a PF formed (second traversal) relative to the traversal in which the PF initially formed. Additionally, metrics describing PF formation itself were calculated. Peak shift was defined as the number of position bins separating the PF formation event from the peak of the post-formation TC, multiplied by cm / bin. A 300-position bin spatial heatmap was used to provide increased spatial resolution. Formation gain was defined as the log_2_ ratio of PF hits on the formation lap divided by the mean post-formation PF hit rate. Peri-formation velocity was defined as the mean animal velocity over the interval [-1.5, 1.0] seconds relative to PF formation. This value was based on the roughly [-4, 2] second asymmetric plasticity kernel observed in BTSP voltage recordings^39, 42, 43^ and designed to capture the densest regions of the kernel distribution, i.e. when most CA1 PN synapses are thought to be potentiated.

##### Predicting properties of newly-formed somatic place fields

Distal tuft dendritic predictors of *de novo* somatic PF properties were selected based on (1) the apparent variability in the timing (relative to somatic activation; ‘lag’) and magnitude (area under the curve; ‘AUC) of tuft activity around the moment of somatic PF formation (Figure 4, E & H; Figure S9) and (2) the observation that lag and AUC did not linearly covary (Figure 4, G & H). Given that these predictors often displayed nonlinear relationships with somatic PF properties (e.g. parabolic and supralinear, see Figure S9), we opted to transform them into polynomial feature space and use these polynomial features to train a double-cross-validated, *L_2_-*regularized (Ridge regularization) linear regression model to predict PF properties. This strategy was chosen over simpler out-of-the-box implementations (e.g. support vector machines or generalized linear models with non-linear kernels) in the interest of accessibility: linear regression weights are relatively straightforward to interpret and implement in various models that may address questions beyond the scope of this study (see table S3 for weights).

Since relationships between tuft features and PF properties varied and could not reasonably be described by a single polynomial order, we transformed each feature into 3^rd^-order polynomial feature space and allowed the Ridge regression model to assign weights to features corresponding to polynomial orders 1-3 for tuft lag and AUC. Poly-transformed features were scaled to [0, 1] prior to model training and testing. Scaled polynomial features were then used to train a leave-one-out double-cross-validated Ridge regression model to predict *de novo* somatic PF properties. Two layers of cross-validation ensured that the model never saw all data points prior to making a prediction and was therefore resistant to overfitting. This approach was used to predict metrics of PF expression and formation described above as well as the point-by-point residuals of the previously reported correlation between an animal’s peri-formation velocity and the width of the resultant somatic PF^39^. Residuals were normalized to their *L_2_* norms in order to account for the fact that larger values tend to display proportionally larger residuals from the least-squares regression line. Scikit-learn^103^ implementations were used for cross-validated Ridge regression (top cross-validation layer was added manually) and polynomial feature space transformation.

#### Statistics

Univariate group comparisons were performed using paired *t*-tests or unpaired *t*-tests with Welch’s correction for heteroscedasticity when both datasets were normally distributed, as determined by D’Agonstino and Pearson test^104^. If at least one dataset was non-normally distributed, then Mann-Whitney *U* tests (unpaired data) or Wilcoxon signed rank tests (paired data) were used. Descriptive statistics of single distributions were treated analogously. Multi-factor group-wise comparisons were performed using ANOVAs or, when missing data points prevented a warranted repeated measures analysis, linear mixed effects models with Geisser-Greenhouse corrections for asphericity. Significant group-wise results were followed up with post-hoc *t*-tests with either Holm-Sidak correction for multiple comparisons or Fisher’s LSD (see table S2 for detailed statistical test descriptions and results). *R* values report Pearson correlation coefficients.

---

**Figure S1.**
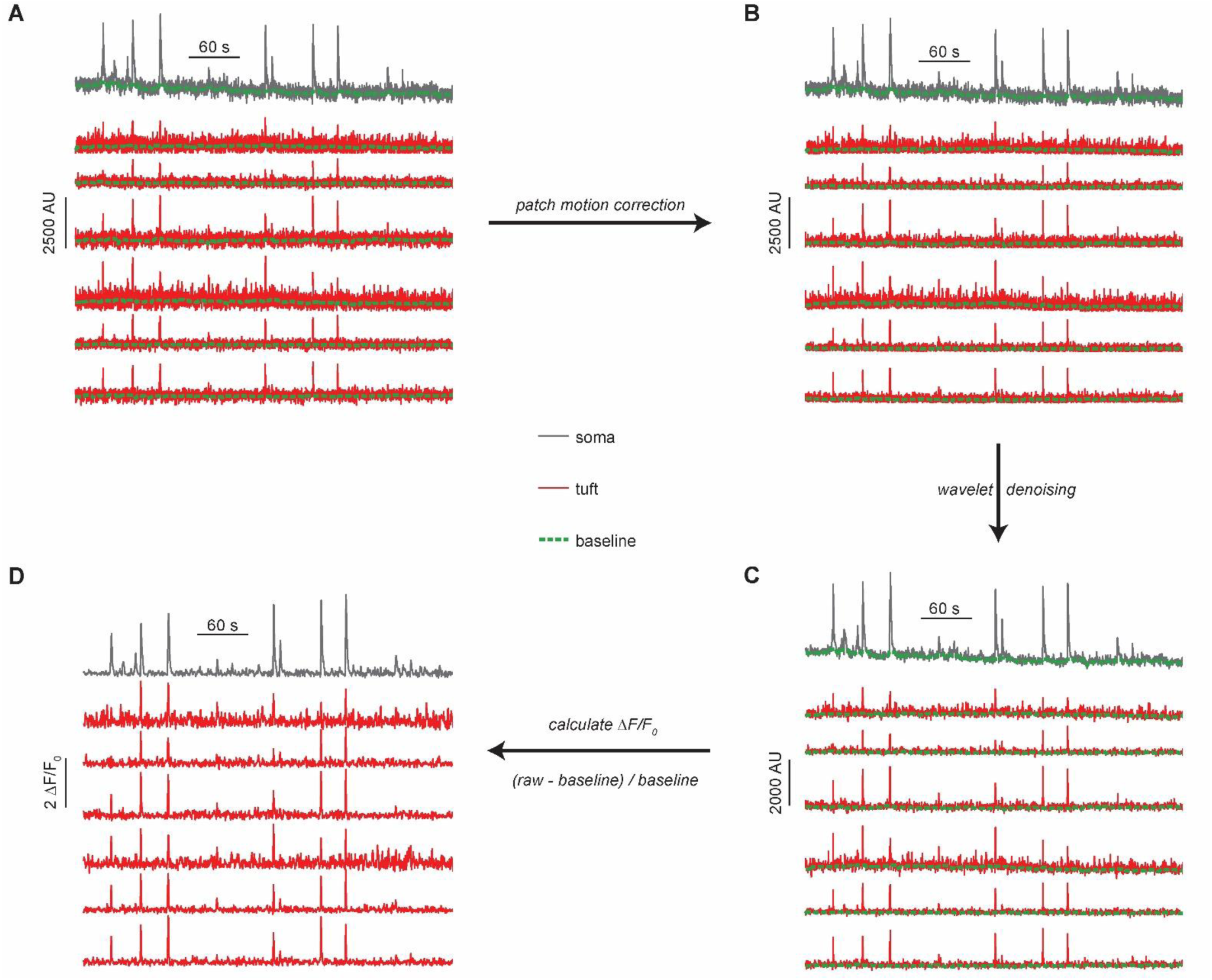
Signal processing. (**A**) Raw fluorescence signals from an example CA1 PN, including the soma (grey) and six distal tuft dendrites (red). Vertical scale is expressed in arbitrary units (AU). Baseline fluorescence (green dashed line) was estimated for each ROI using a rolling maximum of minimum values. (**B**) Raw signals and new baseline estimations after masking frames with uncorrected artifacts due to animal motion. (**C**) Raw signals and final baseline estimations after wavelet-based denoising. (**D**) Final Δ*F/F_0_* used for *Ca^2+^* transient detection and Δ*F/F_0_*-based analyses, calculated from traces and baseline estimates in (C). See supplementary materials and methods for additional detail regarding each processing step. Temporally contiguous subsets of signals are shown, rather than entire 30-min traces, to aid visualization.

**Figure S2.**
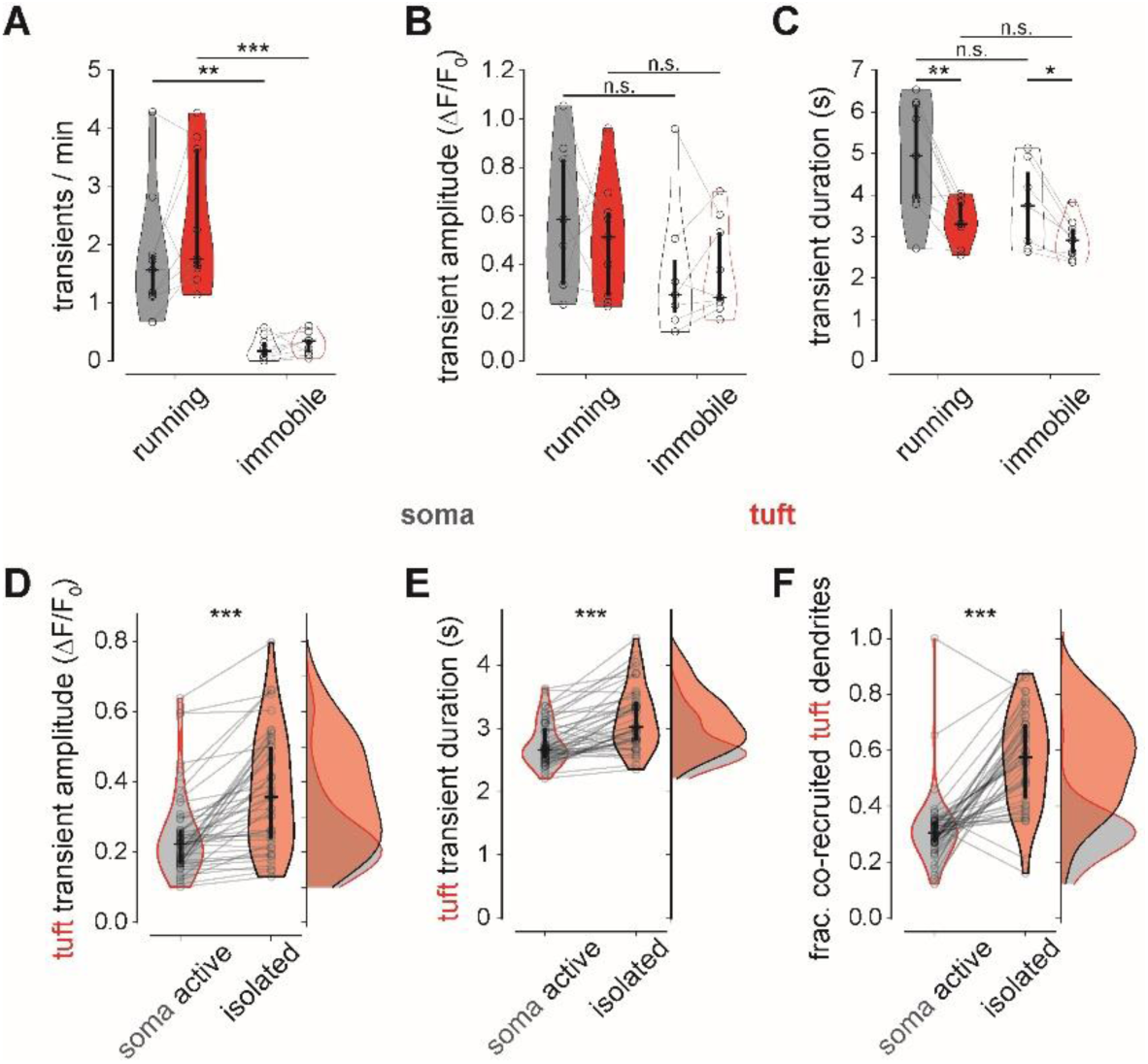
Paired analysis of CA1 PN *Ca^2+^* transient properties by compartment, running state, and cross-compartment recruitment. (**A**) *Ca^2+^* transient frequency for soma (grey) and distal tuft dendrites (red), separated by running (solid) and immobile (open) locomotor states. Data were averaged by cell. Two-way repeated measured ANOVA, compartment effect: *F*_(1,16)_ = 1.18, *P* > 0.05; running effect: *F*_(1,16)_ = 60.80, *P* < 0.001. (**B**) *Ca^2+^* transient amplitude as shown in (A). Linear mixed effects model, compartment effect: *F*_(1,16)_ = 0.27, *P* > 0.05; running effect: *F*_(1,14)_ = 7.54, *P* < 0.05. Note that cross-compartment comparison is not warranted due to confounding disparity in imaging depths. (**C**) *Ca^2+^* transient duration as shown in (A). Linear mixed effects model, compartment effect: *F*_(1,16)_ = 9.96, *P* < 0.01; running effect: *F*_(1,14)_ = 14.86, *P* < 0.01; interaction: *F*_(1,14)_ = 2.0050, *P* > 0.05. (**D**) Amplitude of tuft *Ca^2+^* transients that were accompanied by somatic *Ca^2+^* transients (grey) or isolated (red). Gaussian kernel density estimates shown at right. Data were averaged by dendrite. (**E**) Duration of tuft *Ca^2+^*transients as shown in (D). (**F**) Fraction of imaged tuft dendrites that were co-recruited for a given tuft event, as in (D). Tuft events were defined as epochs containing one or more tuft *Ca^2+^* transients that overlapped in time. Violin plots depict full data range with lines at medians and error bars spanning first to third quartiles. \**P* < 0.05, \*\**P* < 0.01, \*\*\**P* < 0.001. See table S1 for sample sizes and table S2 for additional statistical details.

**Figure S3.**
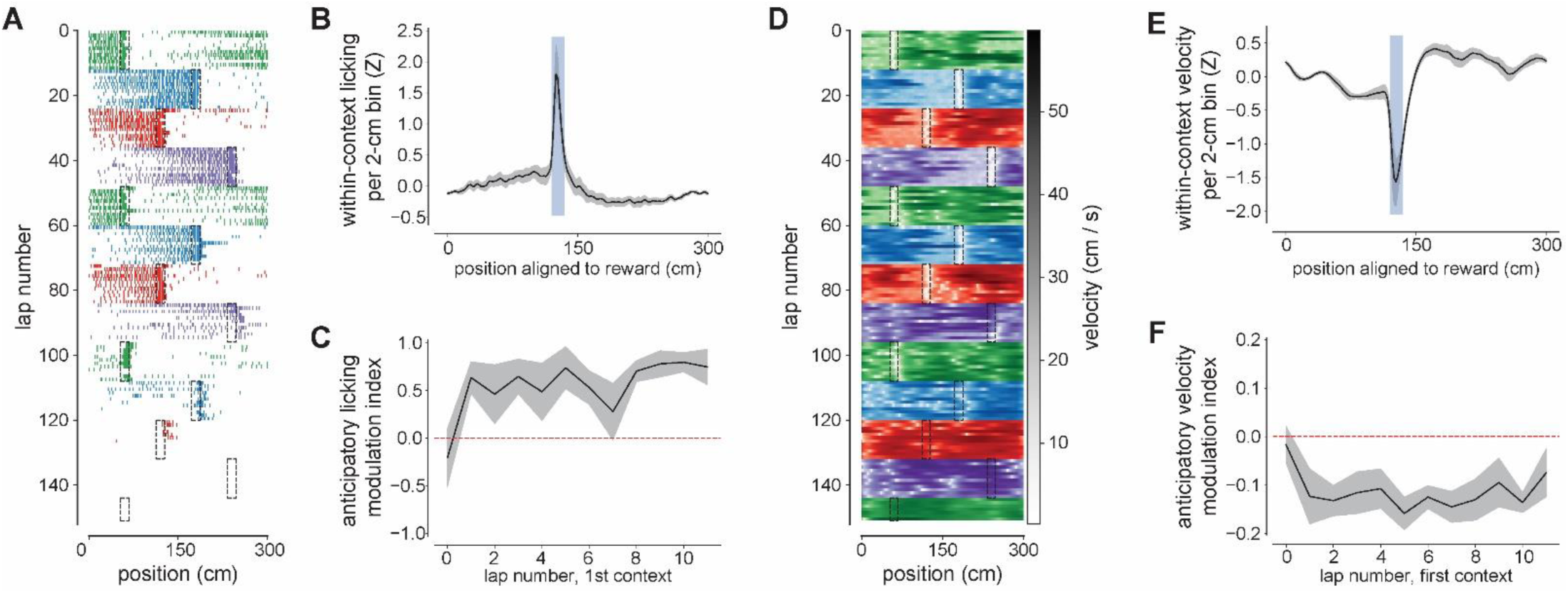
Single-trial learning in a virtual reality-based spatial navigation task. (**A**) Raster plot showing licks as a function of lap number and position along 3-m linear virtual tracks. Ticks are color-coded by contexts. Dashed black rectangles indicate reward zone locations at which water was delivered. (**B**) Licking as a function of position relative to water reward. Lick counts were binned (2 cm), normalized to duration of bin occupancy, and aligned across virtual contexts by reward zone location for each mouse. Blue shaded region indicates water reward zone. (**C**) Anticipatory licking modulation on approach to the reward zone, by lap, in the first exposure to virtual context A (green ticks in panel A). Anticipatory licking was quantified using a modulation index (between -1 and 1) based on the 20% area before reward zone onset (PRE) and the 20% area on the opposite end of the track (BASELINE): (PRE – BASELINE) / (PRE + BASELINE). (**D**) Heatmap of animal velocity analogous to (A). (**E**) Velocity as a function of position relative to water reward as in (B). (**F**) Anticipatory velocity modulation as in (C). Data are shown as mean ± SEM and correspond to the 6 out of 8 imaging sessions that utilized location-fixed reward zones. *N* = 6 mice.

**Figure S4.**
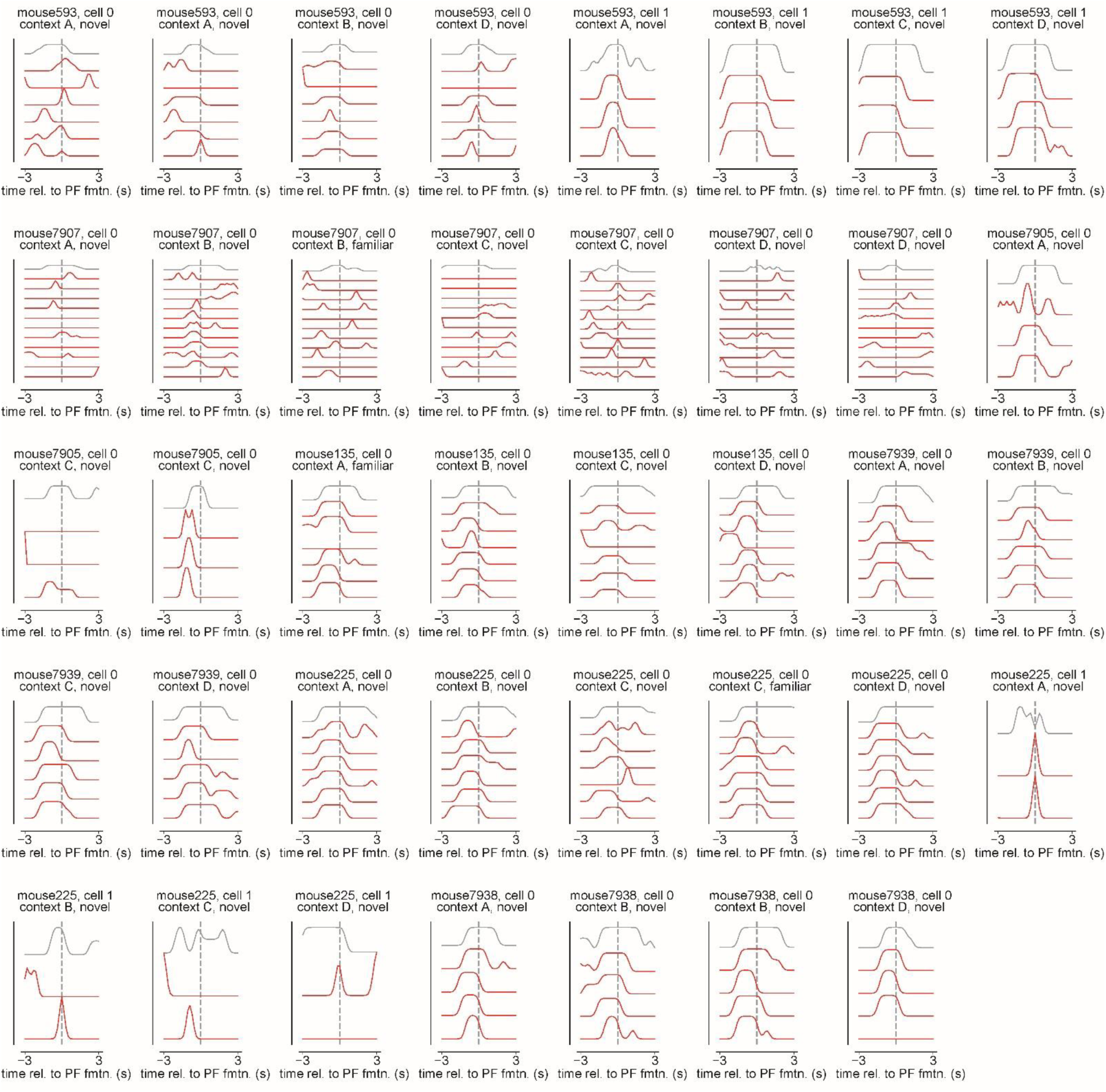
Somatic and tuft activity profiles of all somatic PF formation events. Somatic (grey) and tuft (red) activity profiles for each somatic PF formation event (*N* = 39). Traces represent deconvolved events used to detect PF formation events, smoothed by 1 frame and min-max normalized between 0 and 1. Each panel corresponds to one PF formation event with descriptive title including mouse ID, cell number, context ID, and whether context was novel (initial traversal) or familiar (subsequent traversals). Dashed lines indicate estimated moment of somatic PF formation (see supplementary materials and methods). All traces were resampled to the lowest common frame rate across imaging sessions and smoothed with a 1-frame Gaussian kernel in addition to smoothing used during signal processing (see supplementary materials and methods).

**Figure S5.**
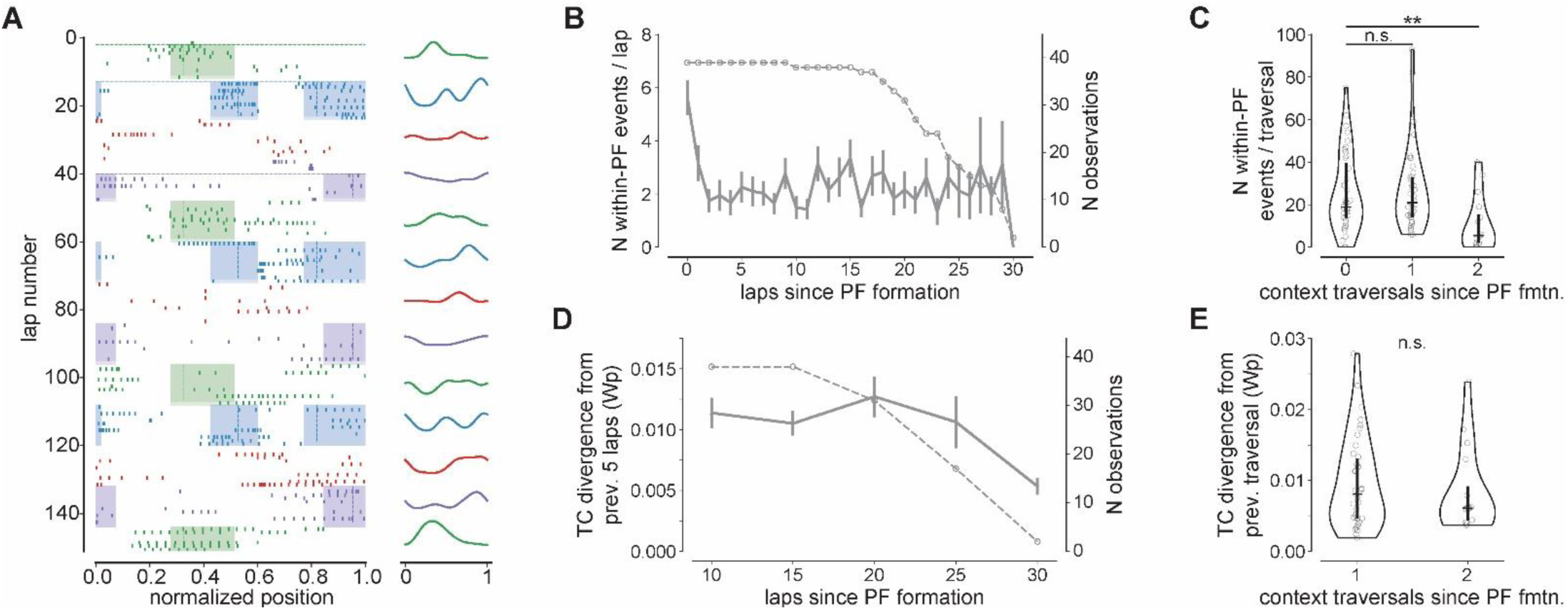
Stability of spontaneously-formed PFs across laps and virtual context traversals. (A) Left: Example raster plot of somatic activity, color-coded by context. Tick marks represent deconvolved events from *ΔF/F_0_* signal which were used for spatial tuning analyses. Each row represents one lap. Shaded regions denote *de novo* PFs, horizontal dashed lines indicate PF formation laps, and vertical dashed lines indicate animal position at the moment of PF formation. Right: Spatial tuning curves (TCs) corresponding to each virtual context traversal, calculated from events shown in raster plot at left. Like-colored TCs illustrate the stability of *de novo* PFs and subthreshold tuning across traversals on a given virtual context. (**B**) Number of events falling within PF bounds as a function of laps since PF formation. Even rates were averaged across contexts for each event. Sample sizes shown on the right y-axis denote the number of PF formation events that were observed across *N* laps within concatenated traversals of the same virtual context. (**C**) Number of events falling within PF bounds as a function of context traversals since PF formation. (**D**) TC divergence, quantified as Wasserstein distance (W_p_), in 5-lap bins relative to PF formation (e.g. lap 10 represents W_p_ between TC in laps 5-9 versus laps 0-4). Data are displayed as in (B). (**E**) TC divergence shown as in (C). Error bars indicate SEM. Violin plots depict full data range with lines at medians and error bars spanning first to third quartiles. \*\**P* < 0.001.

**Figure S6.**
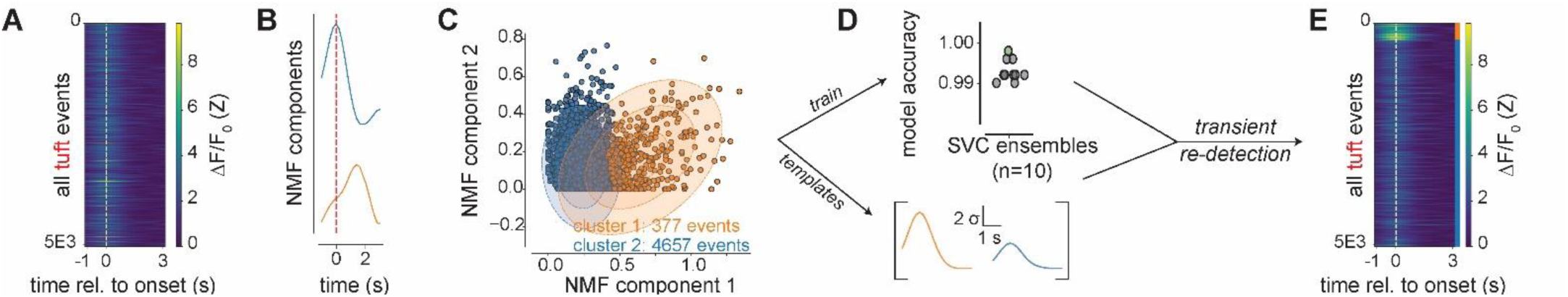
Identification of putative plateau potentials in CA1 PN distal tuft dendrites. (**A** to **C**) Clustering approach to identify putative plateau potentials. Tuft PETHs triggered on *Ca^2+^* transient (*N* = 5,034 from 46 tuft dendrites and 9 CA1 PNs) peaks (A) were factorized into two components (B) and the resulting factorization matrix was clustered using a *K*=2 K-means clustering model (C). See supplementary materials and methods for additional detail. (**D**) Classification approach to label *Ca^2+^* transients as ordinary transients or putative plateau potentials. 10 ensembles of 10 support vector classifiers (SVCs, top) were trained to predict clustering labels and the model with highest accuracy (green fill) was stored. Template-based transient detection was rerun using templates from both clusters (bottom). In cases when an event matched both templates, the trained classifier assigned a ‘transient’ or ‘plateau’ label. (**E**) Final classification of re-detected *Ca^2+^* transients as plotted in (A). Binary color bar at right corresponds to cluster colors in (C) and denotes final label assignment. While clusters inevitably appear poorly separated in (C) due to temporal overlap of NMF components (B), final classifications in fact show clear separation between *Ca^2+^*transients and putative plateaus based on *ΔF/F_0_* signals.

**Figure S7.**
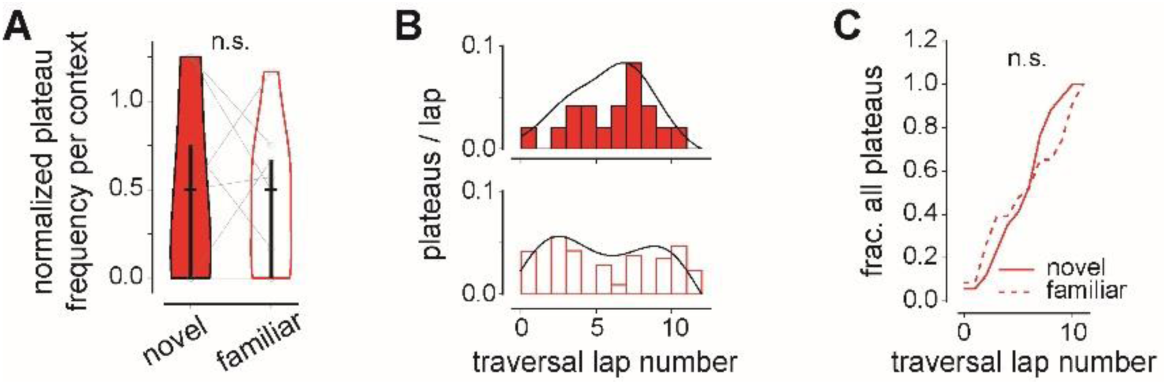
Novel context exposure is not associated with tuft plateau potential prevalence. (**A**) Plateau potential frequency in novel and familiar contexts normalized to number of context traversals (see Figure 3A). Violin bodies span full data range with lines at medians and error bars spanning first to third quartiles. (**B**) Histograms and Gaussian kernel density estimates showing tuft plateau probability per lap within individual novel (top) and familiar (bottom) context exposures. (**C**) Cumulative distribution plot comparing by-lap plateau probability distributions within individual novel (solid line) and familiar (dashed line) context traversals. See table S1 for sample sizes and table S2 for additional statistical details.

**Figure S8.**
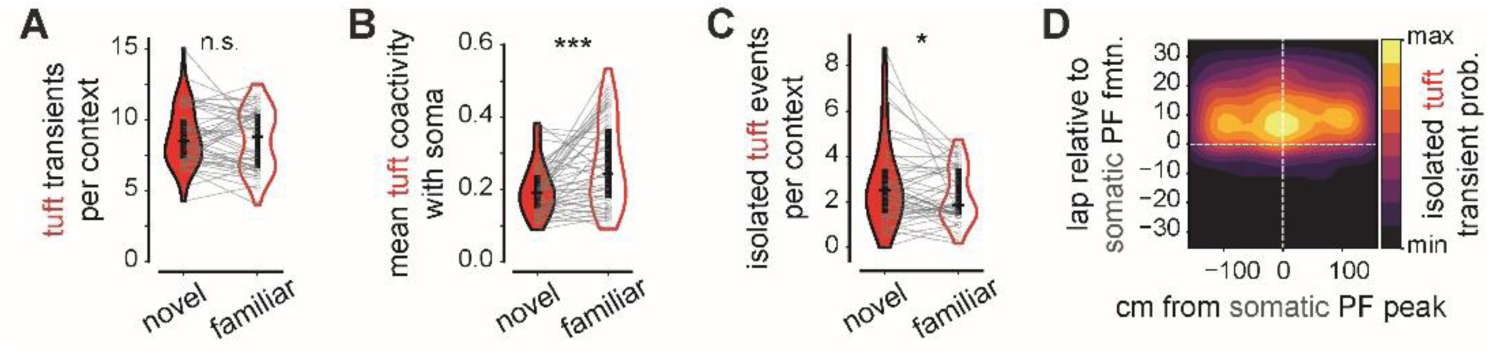
Isolated distal tuft responses to novel context exposure. (**A**) Number of tuft transients per context traversal in novel and familiar contexts. (**B**) Soma-tuft coactivity, i.e. fraction of overlapping transients between soma and a given tuft dendrite, in novel and familiar context traversals. (**C**) Number of isolated tuft transients per context in novel and familiar contexts. (**D**) Heatmap showing the probability distribution of isolated tuft events (2-D Gaussian kernel density estimate) as a function of lap relative to somatic PF formation and distance from somatic PF peak location. Violin bodies span full data range with lines at medians and error bars spanning first to third quartiles. \**P* < 0.05, \*\*\**P* < 0.001. See table S1 for sample sizes and table S2 for additional statistical details.

**Figure S9.**
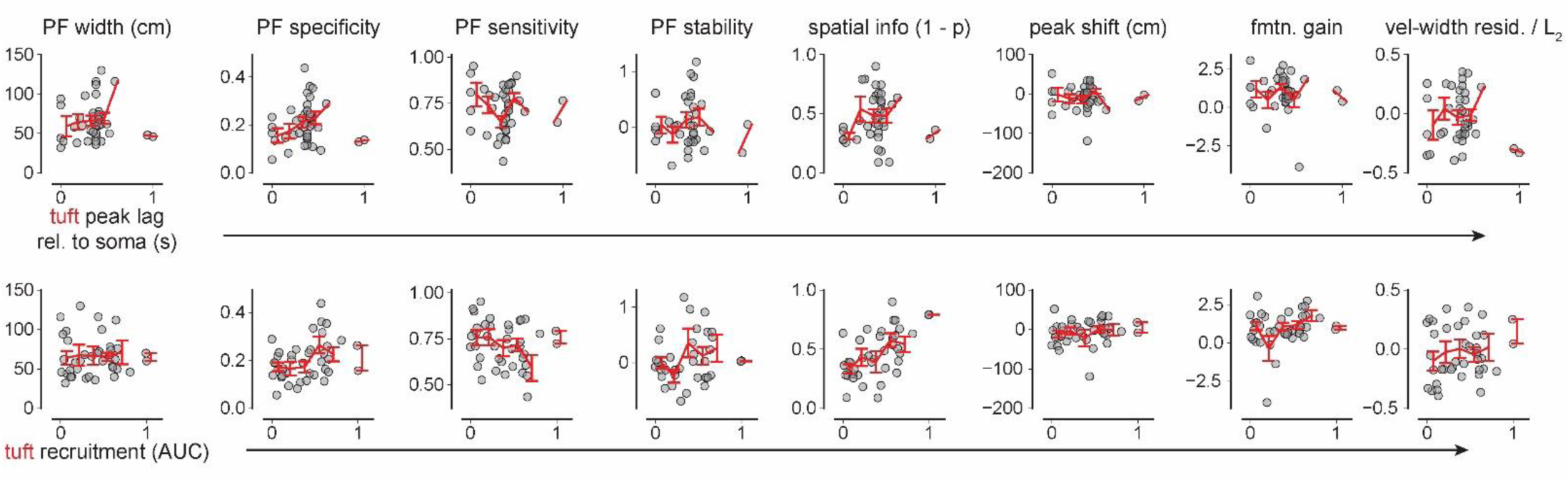
Relationships between somatic PF properties and peri-formation tuft activity features. PF properties (column labels correspond to y-axes) binned according to the time lag between tuft and somatic peri-formation activity profiles (top) and cell-averaged peri-formation tuft response area under the curve (AUC) (bottom). Lag and AUC predictors are min-max scaled from 0 to 1 to show data as they are used to train the regularized linear model described in Figure 5. Red lines and error bars represent mean and SEM of binned PF properties of interest.

**Figure S10.**
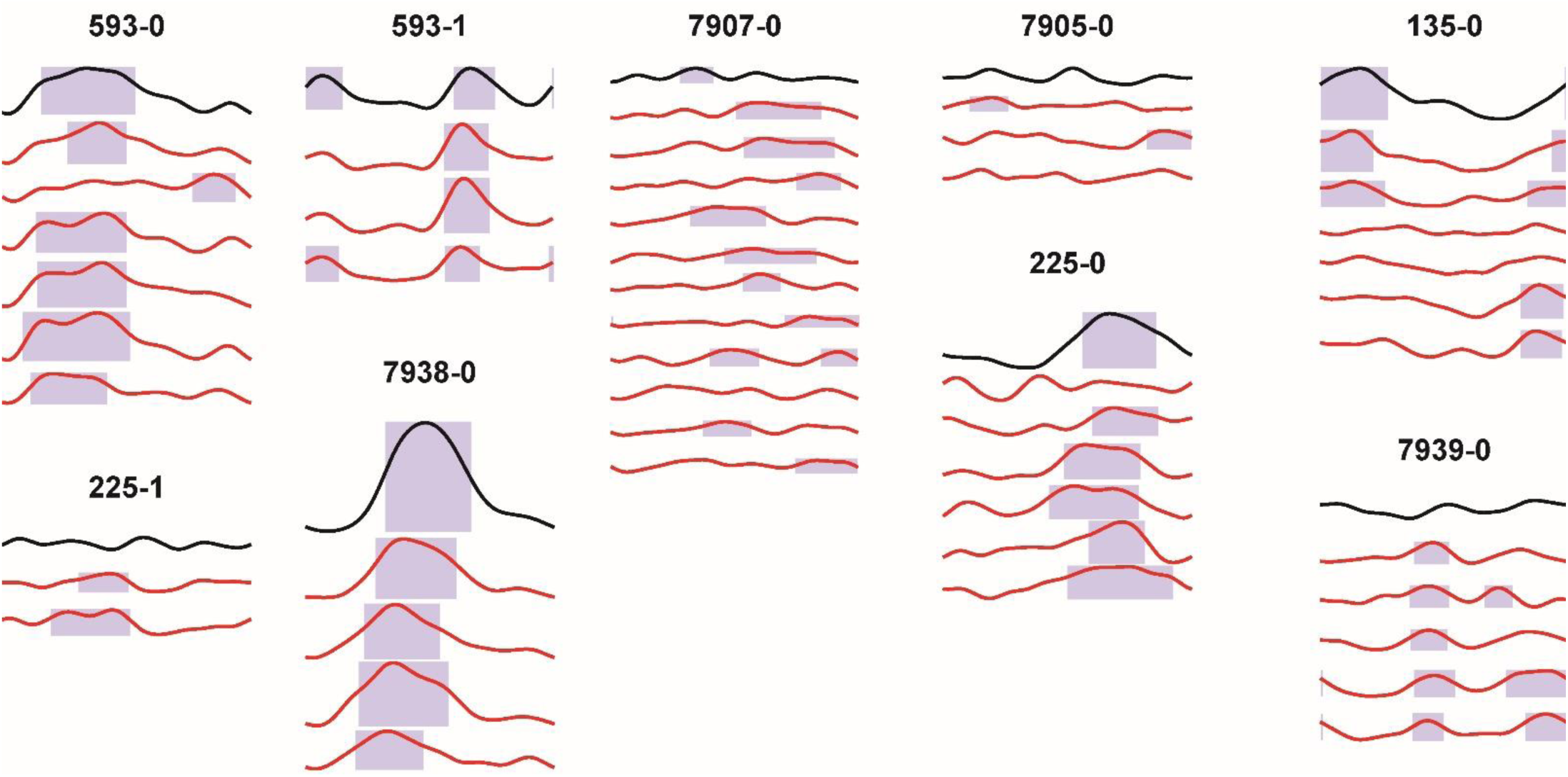
Spatial tuning curves and place fields by cell. TCs and PFs from all imaged regions of interest organized by cell for visual comparison of somatic and tuft dendritic spatial tuning preference. Soma IDs are located above stacked TCs for each cell. Black traces represent somatic TCs and red traces represent tuft TCs. Violet regions indicate statistically significant PFs.

**Figure S11.**
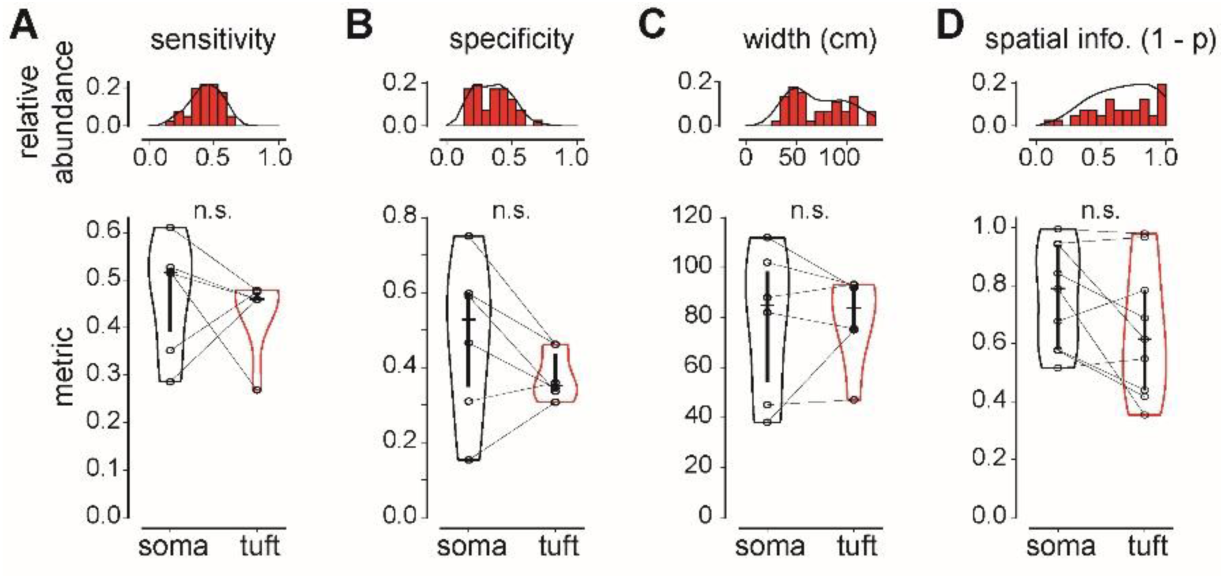
Basic place field properties of CA1 PN distal tuft dendrites. (**A**) PF sensitivity, i.e. the fraction of PF traversals in which at least one event is detected. Top: Histogram and Gaussian kernel density estimate of tuft dendrite PF sensitivity. Bottom: Paired comparison of somatic and cell-averaged tuft PF sensitivity. (**B**) PF specificity, i.e. the fraction of all events falling within PF bounds, as shown in (A). (**C**) PF width as shown in (A). (**D**) Spatial information scores (1 – *P*), as shown in (A), but for all cells independent of PF expression. See supplementary materials and methods for additional detail. Violin plots depict full data range with lines at medians and error bars spanning first to third quartiles. See table S1 for sample sizes and table S2 for additional statistical details.

**Figure S12.**
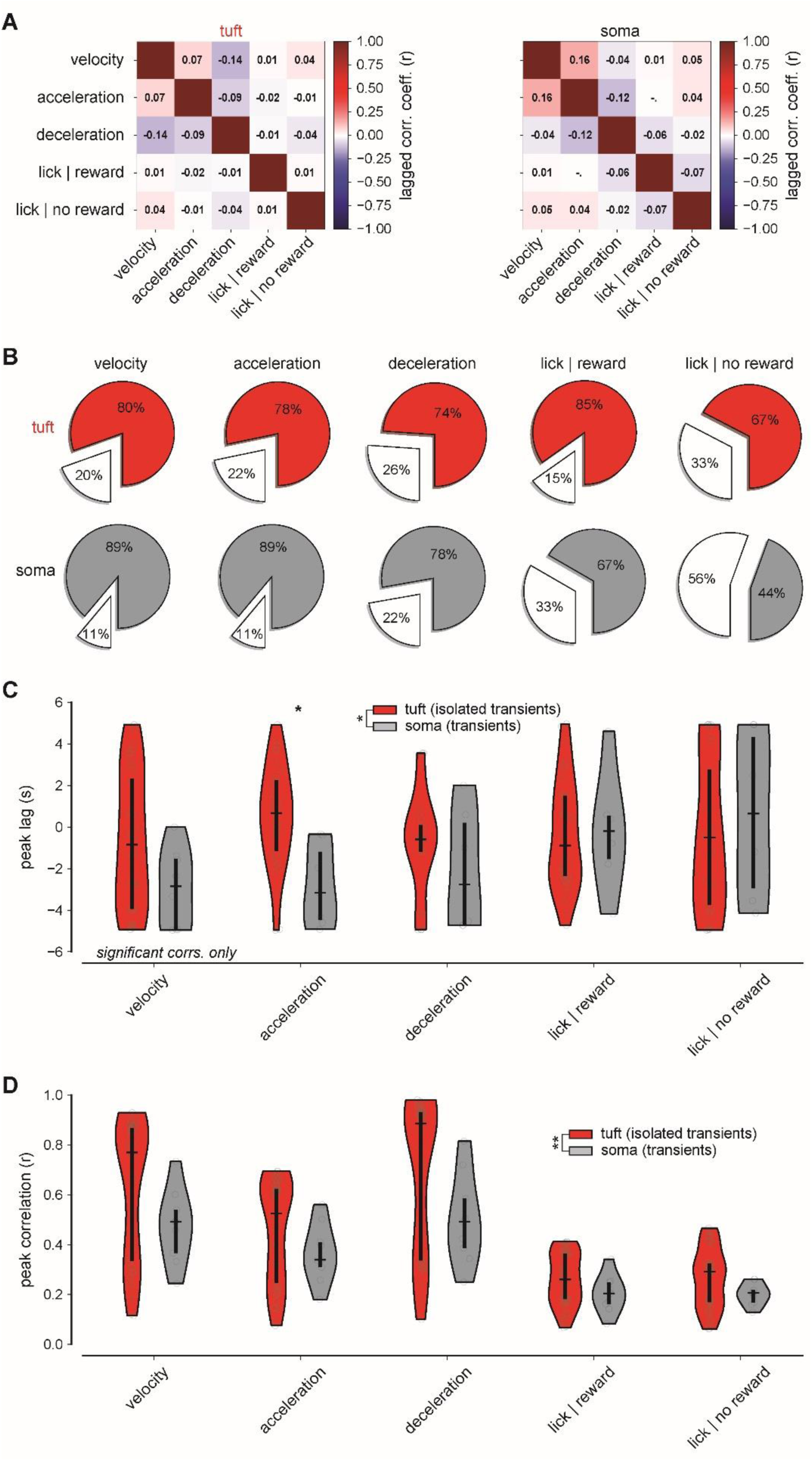
Tuft and somatic non-spatial behavioral representations. (**A**) Correlation matrices showing extent of collinearity (Pearson correlation coefficients) between behavioral features of interest, irrespective of statistical significance. Cross-correlation coefficients were calculated using time lags that resulted in each feature being maximally correlated to isolated tuft *Ca^2+^* transients (left) or somatic *Ca^2+^* transients (right) (see supplementary materials and methods). (**B**) Pie charts showing fractions of tuft dendrites (red, top) and soma (grey, bottom) that were significantly (filled) and non-significantly (open) correlated to behavioral features printed above each column. (**C**) Lags leading to strongest correlations for tuft dendritic (red) and somatic (grey) transient signals across behavioral features denoted in categorical x-tick labels. To accurately portray activity-behavior relationships, lags are only shown for statistically significant correlations. Significance was determined by comparison to distributions of 1,000 null correlation coefficients generated from shuffled data. Two-way ANOVA, compartment effect: *F*_(1, 183)_ = 4.31, *P* < 0.05; behavior effect: *F*_(4, 183)_ = 1.64, *P* > 0.05; interaction: *F*_(4, 183)_ = 2.040, *P* > 0.05. (**D**) Peak correlation coefficients for tuft dendritic and somatic transient signals across behavioral features as shown in (C). All coefficients were included regardless of statistical significance. Two-way ANOVA, compartment effect: *F*_(1, 265)_ = 10.58, *P* < 0.01; behavior effect: *F*_(4, 265)_ =20.89, *P* < 0.001; interaction: *F*_(4, 265)_ = 0.55, *P* > 0.05. Violin plots depict full data range with lines at medians and error bars spanning first to third quartiles. \**P* < 0.05, \*\**P* < 0.01. See table S1 for sample sizes and table S2 for additional statistical details.

**Table S1.**
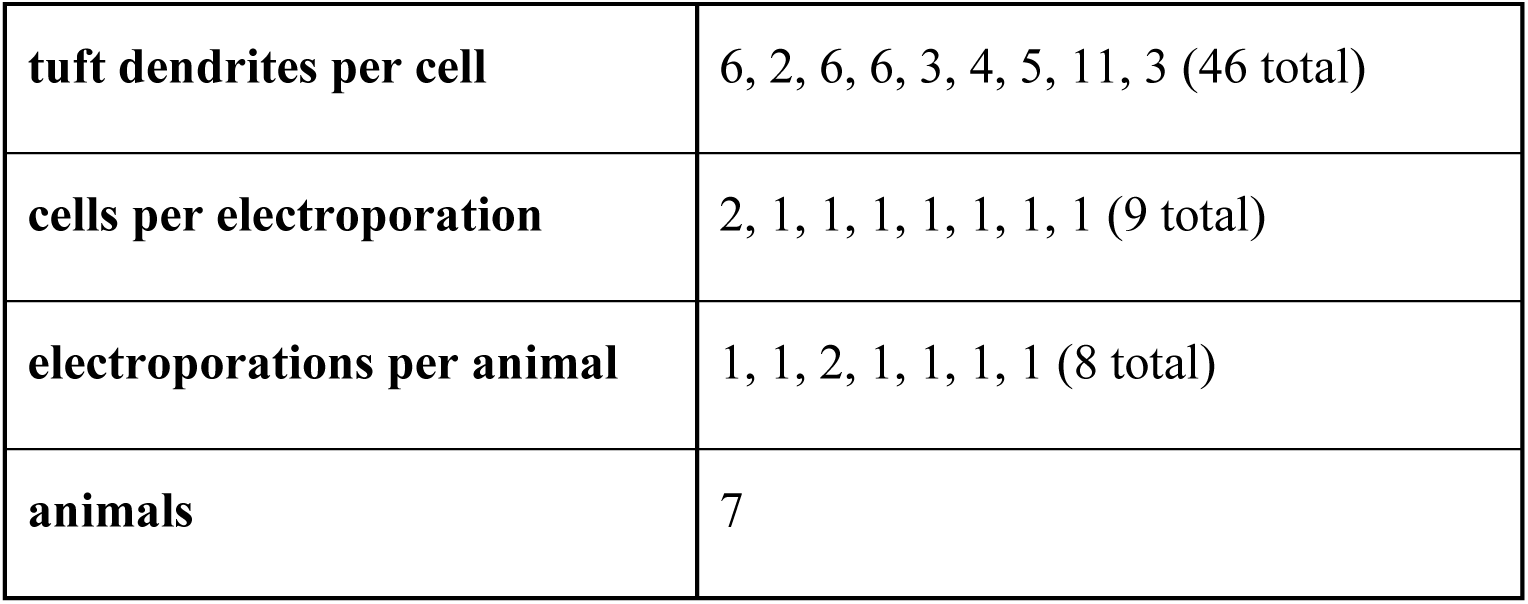
Sample size details. Sample sizes including tuft dendrites per cell, cells per electroporation (i.e. cells per field of view), electroporations per animal (i.e. independent experiments carried out separately in time with new virtual contexts), and number of animals. Values describe single-context ‘random foraging’ experiments corresponding to Figs. 1, 2, & 6 as well as multi-context ‘goal-directed learning’ experiments corresponding to Figs. 3-5.

**Table S2.**
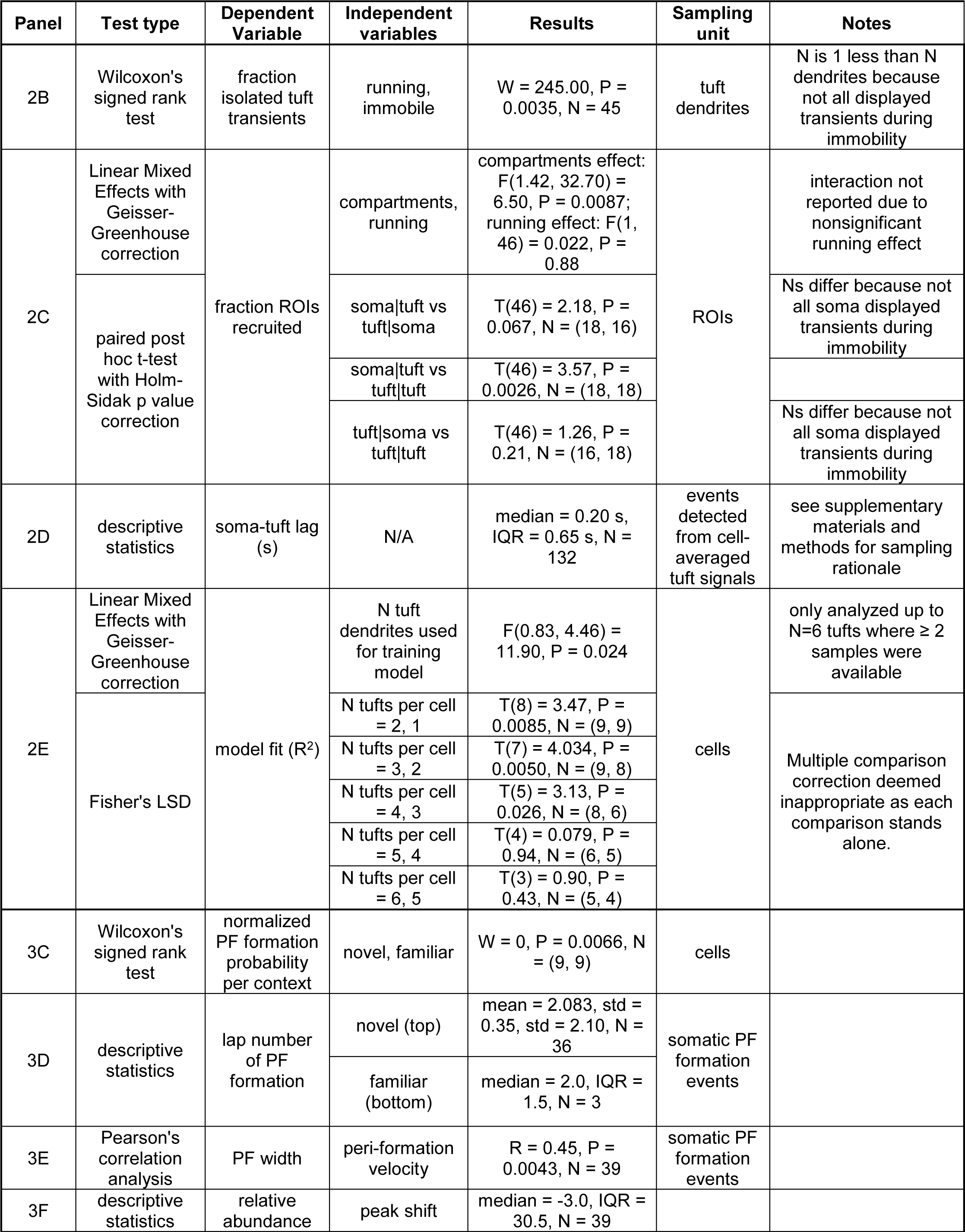
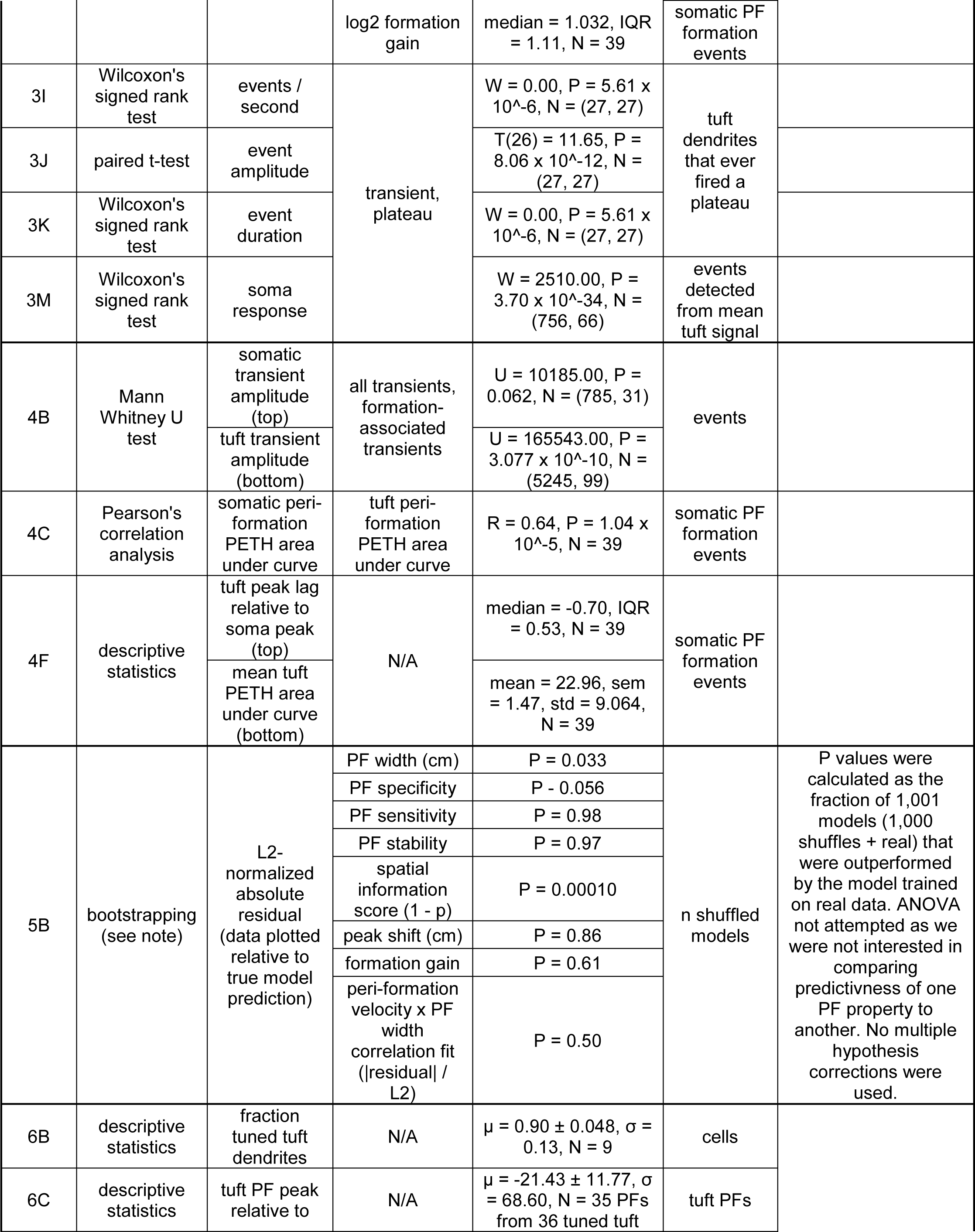
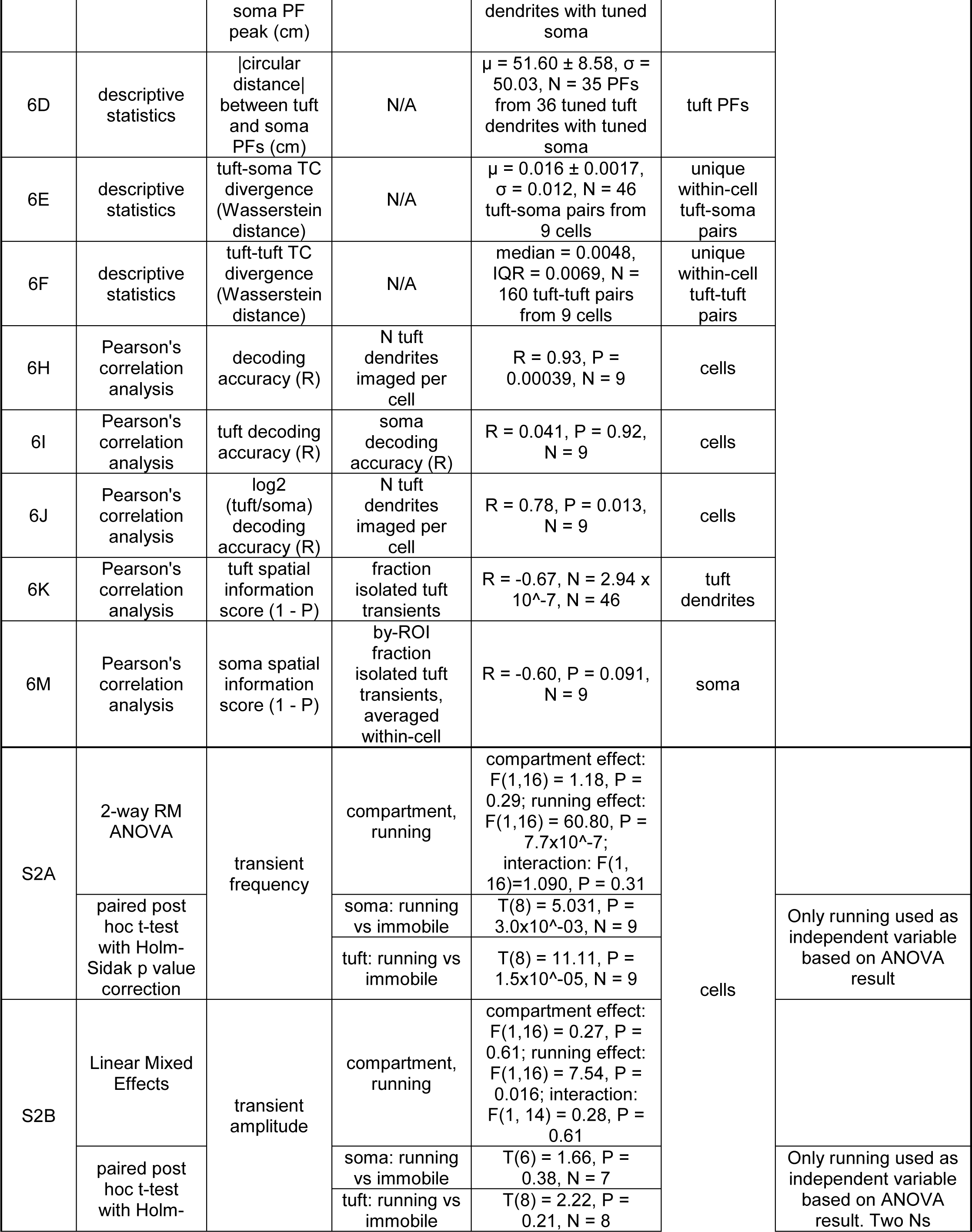
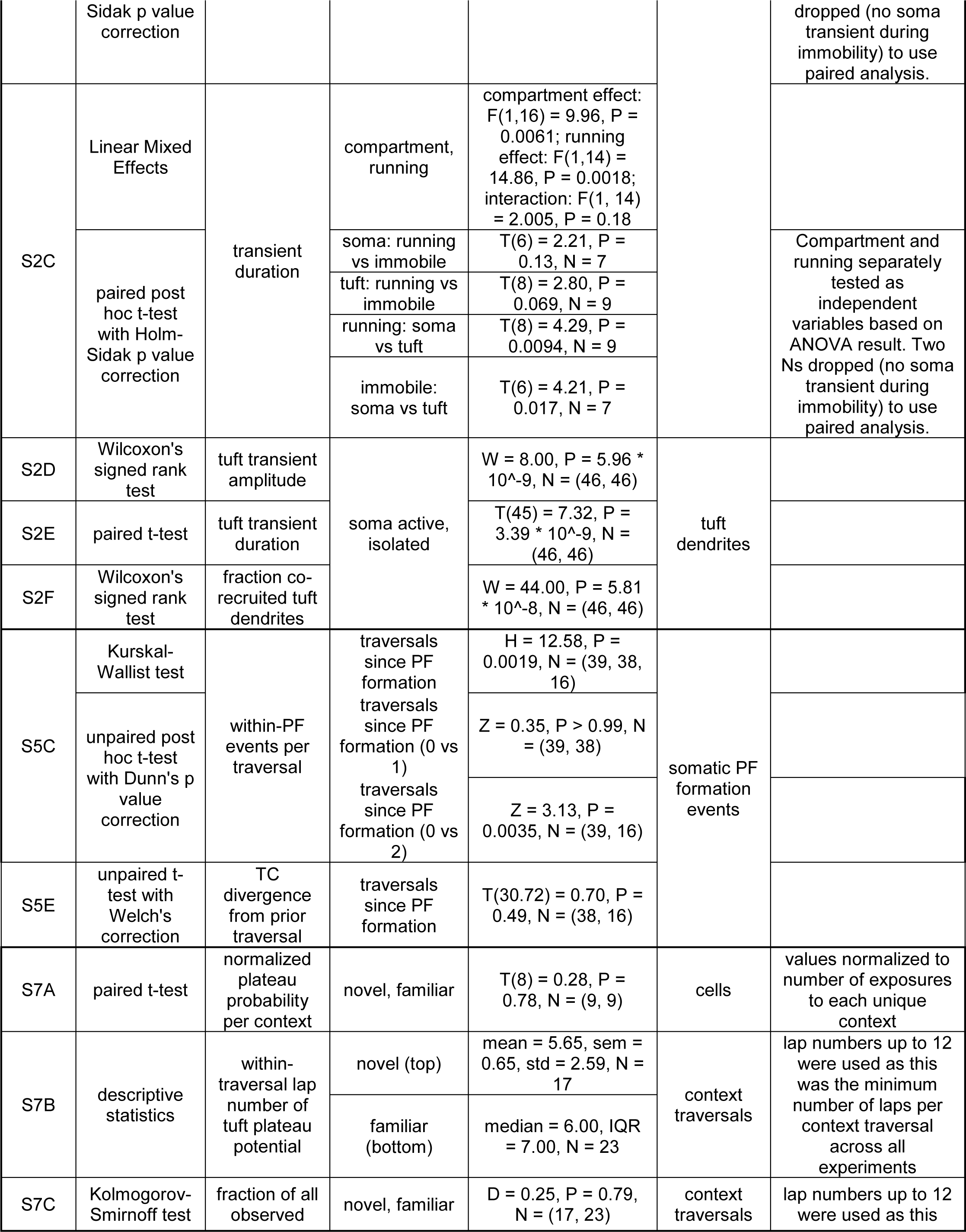
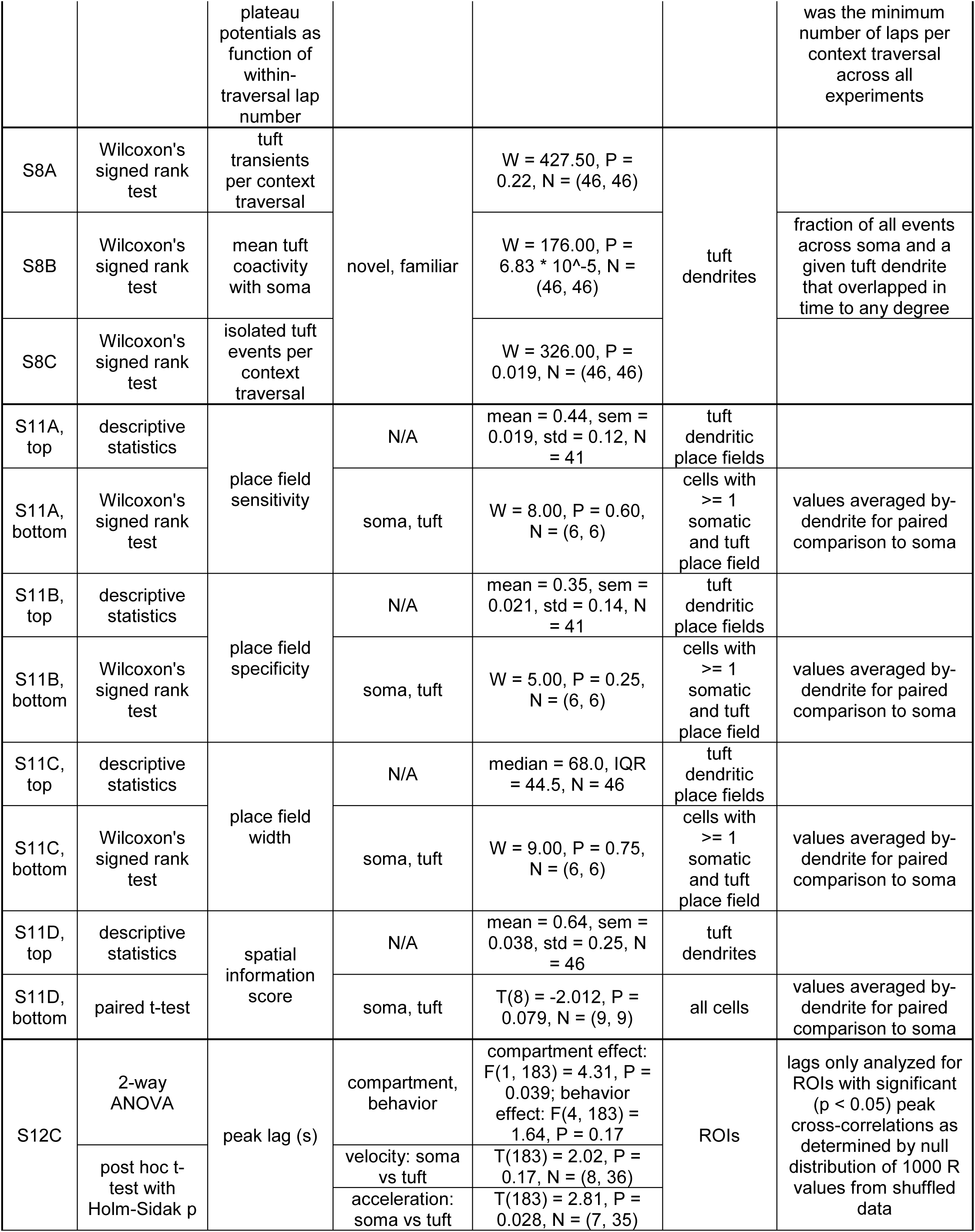
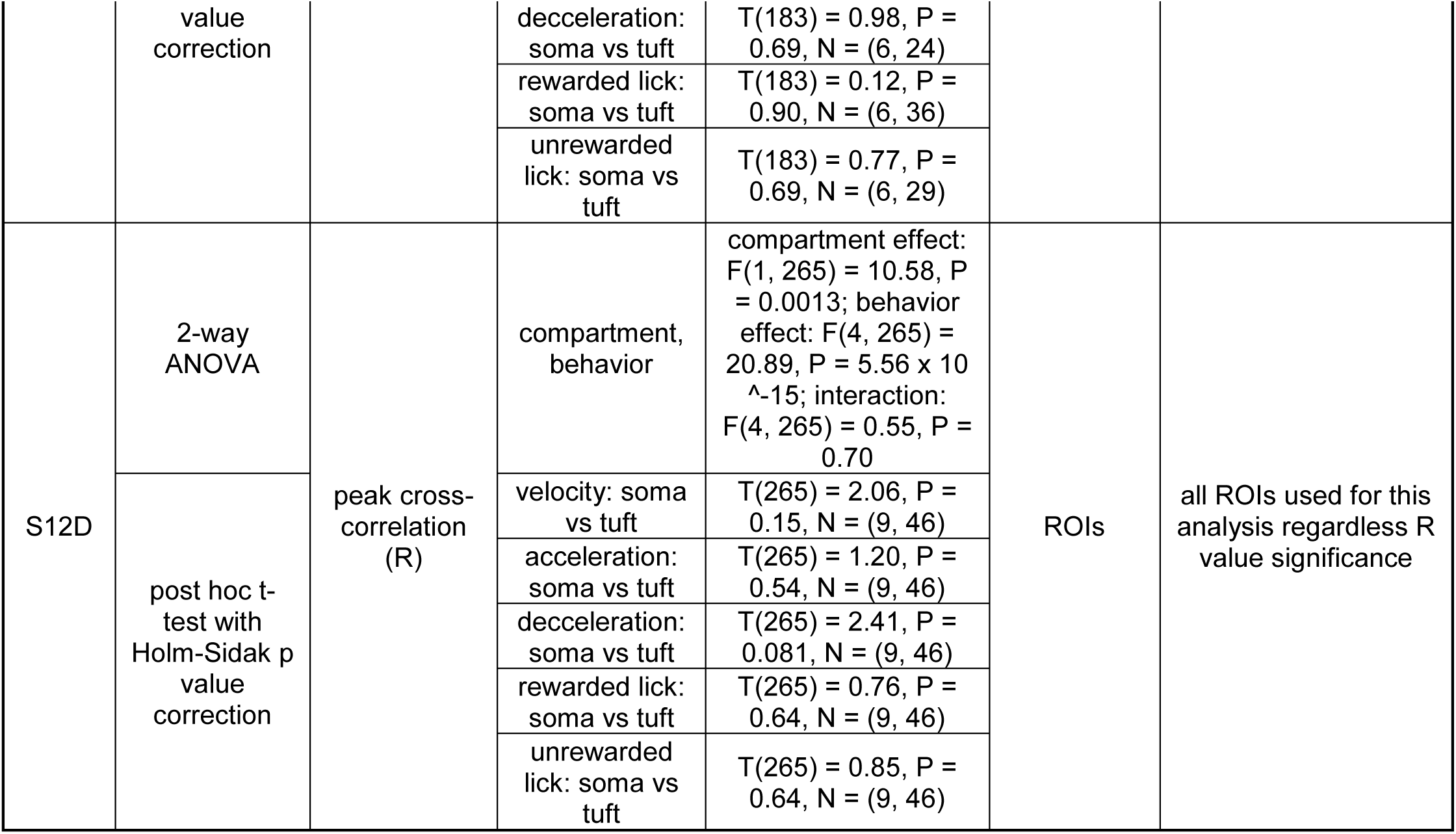
Statistical tests. Results of all group-wise conventional statistical analyses including effect size, degrees of freedom, and *p* values. Non-group-wise analyses, e.g. comparisons against null distributions from shuffled data, are reported in text. See supplementary materials and methods for additional detail.

**Table S3.** Model parameters: predicting PF formation and expression properties based on peri-formation distal tuft activation. Target variable: the PF property to be predicted via double-cross-validated Ridge regression (see supplementary materials and methods). Predictor: the feature of distal tuft activation transformed to 3^rd^ order polynomial space and used to train the Ridge regression model (see Figure 5, A & B). Coeff. a, b, c: Ridge regression weights assigned to each polynomial feature extracted from a given predictor, in order of polynomial degree (1-3).

| target variable | predictor | coeff. a (1°) | coeff. b (2°) | coeff. c (3°) |
| --- | --- | --- | --- | --- |
| width | lag | -4.97 | -22.34 | -0.29 |
| width | AUC | -3.39 | -1.41 | -0.39 |
| specificity | lag | -0.014 | -0.05 | -0.0069 |
| specificity | AUC | 0.047 | 0.029 | 0.0018 |
| sensitivity | lag | 0.0090 | -0.011 | -0.0023 |
| sensitivity | AUC | -0.031 | -0.0066 | 0.0080 |
| stability | lag | 0.068 | -0.10 | 0.028 |
| stability | AUC | 0.074 | 0.041 | 0.0035 |
| spatial information | lag | -0.025 | -0.0043 | -0.017 |
| spatial information | AUC | 0.17 | 0.14 | 0.11 |
| peak shift | lag | 0.40 | -0.031 | 0.64 |
| peak shift | AUC | 2.98 | 3.00 | 2.63 |
| formation gain | lag | -0.0066 | -0.0058 | -0.0016 |
| formation gain | AUC | 0.16 | 0.15 | 0.10 |
| peri-formation velocity-<br>width correlation residual | lag | -0.14 | -0.17 | -0.049 |
| peri-formation velocity-<br>width correlation residual | AUC | -0.015 | 0.018 | 0.054 |

## Notes

### Competing Interest Statement

The authors have declared no competing interest.

### Summary of Updates

Revised strategy to detect place field formation events (stricter validation criteria). Total number of events is 39 compared to previous 46. Updated modeling results (Fig. 5) and statistical reporting (Table S2) stem from this revision. New additions: Fig. S3 (behavioral analyses showing navigational learning), Fig. S5 (Stability of new somatic PFs).

## REFERENCES

1. Elston, G.N., *Cortex,* Cognition and the Cell: New Insights into the Pyramidal Neuron and Prefrontal Function. Cerebral Cortex, 2003. 13(11): p. 1124–1138.

2. Spruston, N., Pyramidal neurons: dendritic structure and synaptic integration. Nature Reviews Neuroscience, 2008. 9(3): p. 206–221.

3. Megıas, M., et al., Total number and distribution of inhibitory and excitatory synapses on hippocampal CA1 pyramidal cells. Neuroscience, 2001. 102(3): p. 527–540.

4. Iascone, D.M., et al., Whole-neuron synaptic mapping reveals spatially precise excitatory/inhibitory balance limiting dendritic and somatic spiking. Neuron, 2020. 106(4): p. 566–578. e8.

5. Dantzker, J. and E. Callaway, Laminar sources of synaptic input to cortical inhibitory interneurons and pyramidal neurons. Nature neuroscience, 2000. 3(7): p. 701–707.

6. Beaulieu-Laroche, L., et al., Enhanced dendritic compartmentalization in human cortical neurons. Cell, 2018. 175(3): p. 643–651. e14.

7. Losonczy, A. and J.C. Magee, Integrative properties of radial oblique dendrites in hippocampal CA1 pyramidal neurons. Neuron, 2006. 50(2): p. 291–307.

8. Larkum, M.E., et al., Synaptic Integration in Tuft Dendrites of Layer 5 Pyramidal Neurons: A New Unifying Principle. Science, 2009. 325(5941): p. 756-760.

9. Smith, S.L., et al., Dendritic spikes enhance stimulus selectivity in cortical neurons in vivo. Nature, 2013. 503(7474): p. 115-120.

10. Polsky, A., B.W. Mel, and J. Schiller, Computational subunits in thin dendrites of pyramidal cells. Nature neuroscience, 2004. 7(6): p. 621–627.

11. Schiller, J., et al., Calcium action potentials restricted to distal apical dendrites of rat neocortical pyramidal neurons. The Journal of physiology, 1997. 505(3): p. 605–616.

12. Major, G., M.E. Larkum, and J. Schiller, Active properties of neocortical pyramidal neuron dendrites. Annual review of neuroscience, 2013. 36: p. 1–24.

13. Xu, N.-l., et al., Nonlinear dendritic integration of sensory and motor input during an active sensing task. Nature, 2012. 492(7428): p. 247-251.

14. Sivyer, B. and S.R. Williams, Direction selectivity is computed by active dendritic integration in retinal ganglion cells. Nature Neuroscience, 2013. 16(12): p. 1848–1856.

15. Jordan, R. and G.B. Keller, Opposing influence of top-down and bottom-up input on excitatory layer 2/3 neurons in mouse primary visual cortex. Neuron, 2020. 108(6): p. 1194–1206. e5.

16. Zhang, S., et al., Organization of long-range inputs and outputs of frontal cortex for top-down control. Nature Neuroscience, 2016. 19(12): p. 1733–1742.

17. Makino, H. and T. Komiyama, Learning enhances the relative impact of top-down processing in the visual cortex. Nature Neuroscience, 2015. 18(8): p. 1116–1122.

18. Losonczy, A., J.K. Makara, and J.C. Magee, Compartmentalized dendritic plasticity and input feature storage in neurons. Nature, 2008. 452(7186): p. 436-441.

19. d’Aquin, S., et al., Compartmentalized dendritic plasticity during associative learning. Science, 2022. 376(6590): p. eabf7052.

20. O’Hare, J.K., et al., Compartment-specific tuning of dendritic feature selectivity by intracellular Ca^2+^ release. Science, 2022. 375(6586): p. eabm1670.

21. Cichon, J. and W.-B. Gan, Branch-specific dendritic Ca^2+^ spikes cause persistent synaptic plasticity. Nature, 2015. 520(7546): p. 180-185.

22. Godenzini, L., A.S. Shai, and L.M. Palmer, *Dendritic Compartmentalization of Learning-Related Plasticity*. eneuro, 2022. 9(3): p. ENEURO.0060-22.2022.

23. Mel, B.W., Synaptic integration in an excitable dendritic tree. Journal of neurophysiology, 1993. 70(3): p. 1086–1101.

24. Poirazi, P. and B.W. Mel, Impact of active dendrites and structural plasticity on the memory capacity of neural tissue. Neuron, 2001. 29(3): p. 779–796.

25. Bicknell, B.A. and M. Häusser, A synaptic learning rule for exploiting nonlinear dendritic computation. Neuron, 2021. 109(24): p. 4001–4017. e10.

26. Tran-Van-Minh, A., et al., Contribution of sublinear and supralinear dendritic integration to neuronal computations. Frontiers in cellular neuroscience, 2015. 9: p. 67.

27. London, M. and M. Häusser, Dendritic computation. Annu. Rev. Neurosci., 2005. 28: p. 503–532.

28. Stuart, G.J. and N. Spruston, Dendritic integration: 60 years of progress. Nat Neurosci, 2015. 18(12): p. 1713–21.

29. Fischer, L., et al., Dendritic Mechanisms for In Vivo Neural Computations and Behavior. Journal of Neuroscience, 2022. 42(45): p. 8460–8467.

30. Zhao, X., et al., Membrane potential dynamics underlying context-dependent sensory responses in the hippocampus. Nature Neuroscience, 2020. 23(7): p. 881–891.

31. Zhao, X., C.-L. Hsu, and N. Spruston, Rapid synaptic plasticity contributes to a learned conjunctive code of position and choice-related information in the hippocampus. Neuron, 2022. 110(1): p. 96–108.e4.

32. O’Keefe, J. and J. Dostrovsky, The hippocampus as a spatial map: preliminary evidence from unit activity in the freely-moving rat. Brain research, 1971.

33. O’keefe, J. and L. Nadel, The hippocampus as a cognitive map. 1978: Oxford: Clarendon Press.

34. Ziv, Y., et al., Long-term dynamics of CA1 hippocampal place codes. Nature neuroscience, 2013. 16(3): p. 264–266.

35. Ferbinteanu, J., P.J. Kennedy, and M.L. Shapiro, Episodic memory—From brain to mind. Hippocampus, 2006. 16(9): p. 691–703.

36. Miller, J.F., et al., Neural activity in human hippocampal formation reveals the spatial context of retrieved memories. Science, 2013. 342(6162): p. 1111-1114.

37. Kolibius, L.D., et al., Hippocampal neurons code individual episodic memories in humans. Nature Human Behaviour, 2023. 7(11): p. 1968–1979.

38. Wixted, J.T., et al., Coding of episodic memory in the human hippocampus. Proceedings of the National Academy of Sciences, 2018. 115(5): p. 1093–1098.

39. Bittner, K.C., et al., Behavioral time scale synaptic plasticity underlies CA1 place fields. Science, 2017. 357(6355): p. 1033-1036.

40. Grienberger, C. and J.C. Magee, Entorhinal cortex directs learning-related changes in CA1 representations. Nature, 2022. 611(7936): p. 554-562.

41. Fan, L.Z., et al., All-optical physiology resolves a synaptic basis for behavioral timescale plasticity. Cell, 2023. 186(3): p. 543–559. e19.

42. Milstein, A.D., et al., Bidirectional synaptic plasticity rapidly modifies hippocampal representations. Elife, 2021. 10: p. e73046.

43. Gonzalez, K.C., et al., Synaptic Basis of Behavioral Timescale Plasticity. bioRxiv, 2023: p. 2023.10. 04.560848.

44. Caya-Bissonnette, L., R. Naud, and J.-C. Béïque, Cellular substrate of eligibility traces. BioRxiv, 2023: p. 2023.06. 29.547097.

45. Xiao, K., et al., A critical role for CaMKII in behavioral timescale synaptic plasticity in hippocampal CA1 pyramidal neurons. bioRxiv, 2023: p. 2023.04.18.537377.

46. Jain, A., et al., Dendritic, delayed, and stochastic CaMKII activation underlies behavioral time scale plasticity in CA1 synapses. BioRxiv, 2023: p. 2023.08. 01.549180.

47. Fyhn, M., et al., Spatial representation in the entorhinal cortex. Science, 2004. 305(5688): p. 1258-1264.

48. Deshmukh, S.S. and J.J. Knierim, Representation of non-spatial and spatial information in the lateral entorhinal cortex. Frontiers in behavioral neuroscience, 2011. 5: p. 69.

49. Giocomo, L.M., et al., Topography of head direction cells in medial entorhinal cortex. Current Biology, 2014. 24(3): p. 252–262.

50. Hardcastle, K., et al., A multiplexed, heterogeneous, and adaptive code for navigation in medial entorhinal cortex. Neuron, 2017. 94(2): p. 375–387. e7.

51. Tsao, A., et al., Integrating time from experience in the lateral entorhinal cortex. Nature, 2018. 561(7721): p. 57-62.

52. Wang, C., et al., Egocentric coding of external items in the lateral entorhinal cortex. Science, 2018. 362(6417): p. 945-949.

53. Høydal, Ø.A., et al., Object-vector coding in the medial entorhinal cortex. Nature, 2019. 568(7752): p. 400-404.

54. Munn, R.G., et al., Entorhinal velocity signals reflect environmental geometry. Nature neuroscience, 2020. 23(2): p. 239–251.

55. Campbell, M.G., et al., Distance-tuned neurons drive specialized path integration calculations in medial entorhinal cortex. Cell reports, 2021. 36(10).

56. Bowler, J.C. and A. Losonczy, Direct cortical inputs to hippocampal area CA1 transmit complementary signals for goal-directed navigation. Neuron, 2023. 111(24): p. 4071–4085. e6.

57. Bittner, K.C., et al., Conjunctive input processing drives feature selectivity in hippocampal CA1 neurons. Nat Neurosci, 2015. 18(8): p. 1133–42.

58. Brun, V.H., et al., Impaired spatial representation in CA1 after lesion of direct input from entorhinal cortex. Neuron, 2008. 57(2): p. 290–302.

59. Remondes, M. and E.M. Schuman, Direct cortical input modulates plasticity and spiking in CA1 pyramidal neurons. Nature, 2002. 416: p. 736.

60. Jarsky, T., et al., Conditional dendritic spike propagation following distal synaptic activation of hippocampal CA1 pyramidal neurons. Nature neuroscience, 2005. 8(12): p. 1667–1676.

61. Takahashi, H. and J.C. Magee, Pathway interactions and synaptic plasticity in the dendritic tuft regions of CA1 pyramidal neurons. Neuron, 2009. 62(1): p. 102–11.

62. Han, E.B. and S.F. Heinemann, Distal dendritic inputs control neuronal activity by heterosynaptic potentiation of proximal inputs. Journal of Neuroscience, 2013. 33(4): p. 1314–1325.

63. Inoue, M., et al., Rational engineering of XCaMPs, a multicolor GECI suite for in vivo imaging of complex brain circuit dynamics. Cell, 2019. 177(5): p. 1346–1360. e24.

64. Kitamura, K., et al., Targeted patch-clamp recordings and single-cell electroporation of unlabeled neurons in vivo. Nature methods, 2008. 5(1): p. 61–67.

65. Wertz, A., et al., Single-cell–initiated monosynaptic tracing reveals layer-specific cortical network modules. Science, 2015. 349(6243): p. 70-74.

66. Rolotti, S.V., et al., Local feedback inhibition tightly controls rapid formation of hippocampal place fields. Neuron, 2022.

67. Bowler, J.C., et al., behaviorMate: An Intranet of Things Approach for Adaptable Control of Behavioral and Navigation-Based Experiments. bioRxiv, 2023: p. 2023.12.04.569989.

68. Rall, W., Theory of physiological properties of dendrites. Annals of the New York Academy of Sciences, 1962. 96(4): p. 1071–1092.

69. Magee, J.C., Dendritic integration of excitatory synaptic input. Nature Reviews Neuroscience, 2000. 1(3): p. 181–190.

70. Magee, J.C., Dendritic Hyperpolarization-Activated Currents Modify the Integrative Properties of Hippocampal CA1 Pyramidal Neurons. The Journal of Neuroscience, 1998. 18(19): p. 7613–7624.

71. Golding, N.L., W.L. Kath, and N. Spruston, Dichotomy of action-potential backpropagation in CA1 pyramidal neuron dendrites. Journal of neurophysiology, 2001. 86(6): p. 2998–3010.

72. Bittner, K.C., B.K. Andrasfalvy, and J.C. Magee, Ion channel gradients in the apical tuft region of CA1 pyramidal neurons. 2012.

73. Bock, T., A. Negrean, and S.A. Siegelbaum, Somatic Depolarization Enhances Hippocampal CA1 Dendritic Spike Propagation and Distal Input-Driven Synaptic Plasticity. The Journal of Neuroscience, 2022. 42(16): p. 3406–3425.

74. Bloss, Erik B., et al., Structured Dendritic Inhibition Supports Branch-Selective Integration in CA1 Pyramidal Cells. Neuron, 2016. 89(5): p. 1016–1030.

75. Chen, K., et al., Persistently modified h-channels after complex febrile seizures convert the seizure-induced enhancement of inhibition to hyperexcitability. Nature Medicine, 2001. 7(3): p. 331–337.

76. Dyhrfjeld-Johnsen, J., et al., Upregulated H-current in hyperexcitable CA1 dendrites after febrile seizures. Frontiers in Cellular Neuroscience, 2008. 2: p. 220.

77. Gasparini, S., M. Migliore, and J.C. Magee, On the initiation and propagation of dendritic spikes in CA1 pyramidal neurons. Journal of Neuroscience, 2004. 24(49): p. 11046–11056.

78. Magee, J.C. and M. Carruth, Dendritic voltage-gated ion channels regulate the action potential firing mode of hippocampal CA1 pyramidal neurons. Journal of neurophysiology, 1999. 82(4): p. 1895–1901.

79. Geiller, T., et al., Local circuit amplification of spatial selectivity in the hippocampus. Nature, 2021: p. 1–5.

80. Sheffield, M.E.J., M.D. Adoff, and D.A. Dombeck, Increased Prevalence of Calcium Transients across the Dendritic Arbor during Place Field Formation. Neuron, 2017. 96(2): p. 490–504 e5.

81. Dong, C., A.D. Madar, and M.E.J. Sheffield, Distinct place cell dynamics in CA1 and CA3 encode experience in new environments. Nature Communications, 2021. 12(1): p. 2977.

82. Priestley, J.B., et al., Signatures of rapid plasticity in hippocampal CA1 representations during novel experiences. Neuron, 2022. 110(12): p. 1978–1992. e6.

83. Golding, N.L., N.P. Staff, and N. Spruston, Dendritic spikes as a mechanism for cooperative long-term potentiation. Nature, 2002. 418(6895): p. 326-331.

84. Hafting, T., et al., Microstructure of a spatial map in the entorhinal cortex. Nature, 2005. 436(7052): p. 801-806.

85. Callaway, J.C. and W.N. Ross, Frequency-dependent propagation of sodium action potentials in dendrites of hippocampal CA1 pyramidal neurons. Journal of neurophysiology, 1995. 74(4): p. 1395–1403.

86. Bloss, E.B., et al., Single excitatory axons form clustered synapses onto CA1 pyramidal cell dendrites. Nature Neuroscience, 2018. 21(3): p. 353–363.

87. Maccaferri, G., et al., Properties of the hyperpolarization-activated current in rat hippocampal CA1 pyramidal cells. Journal of Neurophysiology, 1993. 69(6): p. 2129–2136.

88. Surges, R., T.M. Freiman, and T.J. Feuerstein, Input resistance is voltage dependent due to activation of Ih channels in rat CA1 pyramidal cells. Journal of Neuroscience Research, 2004. 76(4): p. 475–480.

89. Tsay, D., J.T. Dudman, and S.A. Siegelbaum, HCN1 Channels Constrain Synaptically Evoked Ca^2+^ Spikes in Distal Dendrites of CA1 Pyramidal Neurons. Neuron, 2007. 56(6): p. 1076–1089.

90. Spruston, N., et al., Activity-dependent action potential invasion and calcium influx into hippocampal CA1 dendrites. Science, 1995. 268(5208): p. 297-300.

91. Basu, J., et al., Gating of hippocampal activity, plasticity, and memory by entorhinal cortex long-range inhibition. Science, 2016. 351(6269): p. aaa5694.

92. Ascoli, G.A., et al., Local Control of Postinhibitory Rebound Spiking in CA1 Pyramidal Neuron Dendrites. The Journal of Neuroscience, 2010. 30(18): p. 6434–6442.

93. Geiller, T., et al., Large-Scale 3D Two-Photon Imaging of Molecularly Identified CA1 Interneuron Dynamics in Behaving Mice. Neuron, 2020. 108(5): p. 968–983.e9.

94. Klausberger, T., GABAergic interneurons targeting dendrites of pyramidal cells in the CA1 area of the hippocampus. European Journal of Neuroscience, 2009. 30(6): p. 947–957.

95. Leão, R.N., et al., OLM interneurons differentially modulate CA3 and entorhinal inputs to hippocampal CA1 neurons. Nature Neuroscience, 2012. 15(11): p. 1524–1530.

96. Sakalar, E., T. Klausberger, and B. Lasztóczi, Neurogliaform cells dynamically decouple neuronal synchrony between brain areas. Science, 2022. 377(6603): p. 324-328.

97. Hajos, N. and I. Mody, Synaptic communication among hippocampal interneurons: properties of spontaneous IPSCs in morphologically identified cells. Journal of Neuroscience, 1997. 17(21): p. 8427–8442.

98. Cossart, R., et al., Interneurons targeting similar layers receive synaptic inputs with similar kinetics. Hippocampus, 2006. 16(4): p. 408–420.

99. Klausberger, T., et al., Complementary roles of cholecystokinin-and parvalbumin-expressing GABAergic neurons in hippocampal network oscillations. Journal of Neuroscience, 2005. 25(42): p. 9782–9793.

100. Cone, I. and H.Z. Shouval, Behavioral time scale plasticity of place fields: mathematical analysis. Frontiers in computational neuroscience, 2021. 15: p. 640235.

101. Longair, M.H., D.A. Baker, and J.D. Armstrong, Simple Neurite Tracer: open source software for reconstruction, visualization and analysis of neuronal processes. Bioinformatics, 2011. 27(17): p. 2453–2454.

102. Friedrich, J. and L. Paninski, Fast active set methods for online spike inference from calcium imaging. Advances In Neural Information Processing Systems, 2016. 29: p. 1984–1992.

103. Pedregosa, F., et al., Scikit-learn: Machine learning in Python. the Journal of machine Learning research, 2011. 12: p. 2825–2830.

104. D’agostino, R. and E.S. Pearson, Tests for departure from normality. Empirical results for the distributions of b 2 and√ b. Biometrika, 1973. 60(3): p. 613-622.

